# Structural basis of caveolin-driven membrane bending

**DOI:** 10.64898/2026.02.05.703862

**Authors:** Sarah M. Connolly, Leon Bergner, Ajit Tiwari, Tyler S. Brant, Samantha Medford, Siddanth Ramesh, Elizabeth D. Tidwell, Youngki Yoo, Ke Xiao, J Gentry, Louise Chang, Bing Han, Padmini Rangamani, Milka Doktorova, Anne K. Kenworthy, Shyamal Mosalaganti, Melanie D. Ohi

## Abstract

Caveolins are monotopic membrane proteins essential for caveolae formation and have key roles in signaling and lipid regulation. Caveolins assemble into amphipathic discs with a central β-barrel, an architecture distinct from other membrane-remodeling proteins. These discs embed in the membrane inducing membrane curvature. However, the mechanism of disc-driven bending remains unresolved. Using cryo-electron tomography, structure-guided mutagenesis, and mammalian cell studies, we show that evolutionarily distinct caveolins differ dramatically in their ability to curve membranes despite their conserved architecture. Through computational and theoretical analyses, we demonstrate that patterning of hydrophobic residues along the rim of the disc of human Caveolin-1 induces the deformation of the surrounding leaflet, dictating membrane bending. Finally, we determine a 4.1Å resolution structure of Caveolin-1 within heterologous caveolae *in situ,* showing the disc adopts a funnel-like conformation, further shaping membrane architecture. Together, these findings reveal fundamental structural principles that empower caveolins to sculpt and remodel cellular membranes.

## Introduction

Cell membranes are dynamic molecular assemblies composed of phospholipids that form bilayers interspersed with cholesterol and membrane proteins. Membranes establish the physical boundary of the cell and allow segregation of intracellular compartments. The shapes of cellular membranes are tightly regulated to adapt to changing physiological demands ^1,2^. These regulated shape transformations rely on proteins that interact with the lipid bilayer to change its architecture and generate curvature. Proteins can remodel membrane morphology through several mechanisms, including inserting amphipathic helices into the bilayer, creating scaffolds that organize lipids into curved arrangements, or producing asymmetric forces between the two leaflets of the bilayer ^3–10^. They can also stabilize membrane structure through oligomerization, anchor the bilayer to the cytoskeleton, and preferentially interact with regions of specific curvature ^3,9–12^. Examples of proteins that drive changes in membrane curvature include clathrin and members of dynamin family, which coordinate vesicle budding at the plasma membrane ^13–16^; Bin/Amphiphysin/Rvs (BAR) domain proteins, which sculpt membranes into tubules and invaginations ^17–19^; the endosomal sorting complex required for transport (ESCRT) machinery, which remodels membranes during intracellular cargo sorting ^20–22^; and caveolins, which form caveolae, 50-100 nm flask-shaped invaginations in the plasma membranes of many vertebrate cells ^23–27^.

Caveolins are monotopic integral membrane proteins that remain constitutively associated with the bilayer, making them unique among curvature-generating proteins. Caveolin forms oligomeric 8S complexes that are believed to drive curvature, establish membrane microdomains, and interact dynamically with other caveolae-associated proteins, most notably the cavins ^25,28–30^. Caveolae serve as central hubs for integrating cellular responses to changes in membrane tension and stress, in part due to their ability to reversibly flatten and disassemble ^30–36^. Disruption of caveolin function, whether through altered expression or mutations, impairs caveolae formation and underlies a broad spectrum of human diseases, including cardiovascular disorders, cancers, and metabolic syndromes, highlighting their vital contribution to cellular homeostasis ^26,34,37–41^. Critically, the molecular basis of caveolin’s role in caveolae biogenesis and regulation remains unclear.

The absence of high-resolution structures of any caveolin for many decades meant that their function in caveolae biogenesis was inferred mainly from studies using mutant or truncated proteins. These analyses identified regions within the proteins that are critical for oligomerization (oligomerization domain, OD) and both membrane attachment and protein-protein interactions (caveolin scaffolding domain, CSD) ^42–47^ (Figure S1A). It also contains a hydrophobic region (intramembrane domain, IMD) that is proposed to form an α-helical hairpin loop in the membrane ^45,48–52^ (Figure S1A). These observations led to the prevailing model that, in the context of full-length caveolin, the α-helical IMD hairpin would insert into the cytoplasmic leaflet of the lipid bilayer, forming a wedge that induces membrane curvature ^10,48,49,53–55^ (Figure S1B).

In contradiction, recent structural analyses of human (*Hs*), purple sea urchin (*Strongylocentrotus purpuratus, Sp*), and choanoflagellate (*Salpingoeca rosetta, Sr)* Caveolins using single particle cryo-electron microscopy (cryo-EM), as well as AlphaFold predictions of additional caveolins, show that the oligomeric complexes organize into amphipathic discs with a central hydrophobic β-barrel ^56,57^. The flat faces of the discs are formed by tightly packed spiraling α-helices, with the periphery of the disc composed of a thicker outer rim. Interestingly, the regions of caveolin identified by mutational and deletion analyses do not match the structural features observed in the cryo-EM maps ^56,57^ (Figure S1C, D). For example, the IMD residues are part of a continuous α-helical region that contributes to the formation of the outer rim and flat membrane-facing surface of the complex rather than forming a discrete hairpin-like structure as predicted in earlier models. These findings suggest an alternative model for how caveolins bend membranes and drive caveolae biogenesis. In this model, the disc embeds into one leaflet of the bilayer, displacing lipids and inducing asymmetric leaflet remodeling that drives curvature (Figure S1E). For the *Hs* Caveolin-1 (*Hs* CAV1) complex, *in vitro* studies using Langmuir monolayers and lipid vesicles demonstrated that the disc is indeed embedded directly into one membrane leaflet ^58^. Complementary molecular dynamics (MD) simulations predict that the *Hs* CAV1 disc stably integrates into the cytosolic leaflet by displacing lipids and inducing localized membrane deformation, which could promote large-scale bending ^58–61^. The general disc-shaped architecture of caveolin oligomers is shared across evolutionarily distant caveolins, suggesting that a similar membrane-remodeling strategy may be exploited throughout Metazoa ^57^.

Together, the structural and modeling studies predict that caveolin discs function as modular building blocks that tile the surface of the caveolar bulb. While several models have been proposed to explain how flat disc-shaped complexes could precisely sculpt membranes to form caveolae ^62,63^, these models remain untested experimentally. Which conserved structural features of caveolin complexes are essential for this process, and whether these mechanisms are conserved across evolutionarily distinct caveolins, are also unknown.

Here, using a combination of *in situ* cryo-electron tomography (cryo-ET), single-particle cryo-EM, cellular microscopy, MD simulations, and modeling, we dissect the structural determinants of caveolin complexes required for membrane bending. We show that despite sharing a similar structural framework with the *Hs* CAV1 complex, the evolutionarily distinct *Sp* and *Sr* Caveolins neither sculpt membranes nor induce caveolae biogenesis. Through mutational analysis and domain swaps, we identify residues located at the edge of the caveolin discs as essential for membrane sculpting. MD simulations highlight the functional significance of these residues by linking the hydrophobicity profile of the complex to the deformation of the surrounding membrane leaflet. These results provide a molecular basis for why *Sp* and *Sr* Caveolin complexes fail to induce membrane curvature despite sharing a conserved global architecture with *Hs* CAV1. Finally, we show that *Hs* CAV1 adopts a funnel-shaped conformation *in situ,* in contrast to the flat conformation of the disc observed in the detergent-solubilized micelles *in vitro*. This altered conformation helps distort the leaflets’ morphology, thereby reshaping the lipid bilayer architecture.

## Results

### The conserved disc architecture of caveolins is insufficient for membrane bending

Recent advances have yielded sub-3.5 Å cryo-EM structures of *Hs* CAV1, *Sp* Caveolin, and *Sr* Caveolin, providing molecular insight into caveolin architecture across evolutionarily distant species ^56,57^ (Figure 1A-C). Despite low sequence identity between *Hs* CAV1 and *Sp* (13%) or *Sr* Caveolin (16%), all three assemble into conserved complexes comprising 11 protomers arranged symmetrically as an amphipathic disc. Within each complex, several structural elements are conserved, including an N-terminal variable region, a hook structure, a scaffolding domain (SD), a spoke region (SR), a β-strand, and, unique to *Sp* and *Sr* Caveolins, a C-terminal variable region (Figure 1A-C and Figure S1C, D). While the overall architecture of these complexes is similar, key differences include disc diameter, curvature variation, and β-barrel length (Figure 1A-C). To assess whether the conserved disc-shaped structure of caveolin complexes is sufficient to drive membrane bending, we tested *Sp* and *Sr* Caveolins in an established prokaryotic model. Expressing *Hs* CAV1 in *Escherichia coli (E. coli)* causes the inner membrane to form vesicles known as heterologous (*h*-) caveolae, which are enriched in caveolins and resemble mammalian caveolae in size (∼50-100 nm) ^24,64^. Mutant caveolins that cannot oligomerize or generate caveolae in mammalian cells also fail to produce *h*-caveolae ^24,64^. At the same time, this simplified model bypasses the requirements for specific lipids and additional proteins needed to build caveolae in vertebrates ^30^. Thus, this system recapitulates key features of mammalian caveolin function, including oligomerization and membrane shaping, and serves as an effective model for studying caveolin-mediated membrane bending.

**Figure 1.**
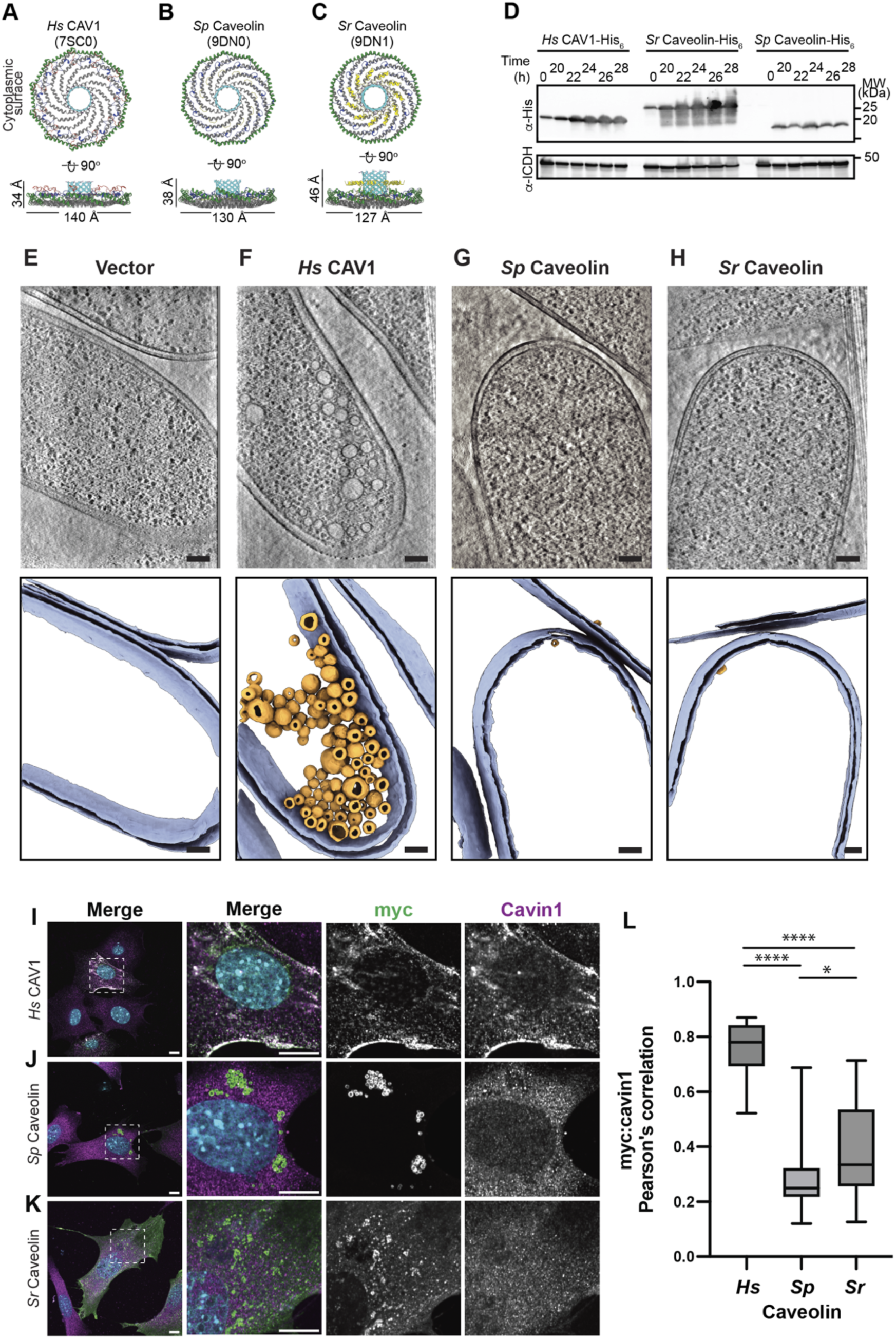
*S. purpuratus (Sp)* and *S. rosetta (Sr)* Caveolins do not sculpt membranes or rescue caveolae biogenesis. (**A-C**) Models from cryo-EM structures of Human (*Hs*), *S. purpuratus* (*Sp*), and *S. rosetta* (*Sr*) Caveolin complexes. Cytoplasmic and side views are shown. Domain coloring: N-terminal variable region (yellow), pin motif (red), scaffolding domain (green), spoke region (gray), β-strand (cyan). (**D**) Western blot analysis showing expression of His_6_-tagged *Hs*, *Sp*, or *Sr* Caveolins in *E. coli* over 28 hours. Samples were harvested and vitrified for cryo-ET analysis at 24 hours. The membrane was probed with anti-His_6_ (α-His) or anti-ICDH (α-ICDH) antibodies. (**E-H**) Cryo-ET analysis of *E. coli* expressing His_6_ vector (E), *Hs* CAV1-His_6_ (F), *Sp* Caveolin-His_6_ (G), or *Sr* Caveolin-His_6_ (H). Slices through a representative deconvoluted tomogram (top panel) and the corresponding segmentation (bottom panel). Bacterial membranes are shown in blue, and *h*-caveolae are shown in gold. Scale bars, 100 nm. **(I-K**) Representative confocal microscopy images shown for CAV1^-/-^ MEF cells expressing myc-tagged *Hs* CAV1, *Sp* Caveolin, or *Sr* Caveolin. Transfected cells expressing indicated constructs for 24 h were fixed and immunostained for myc (green) and Cavin1 (magenta). Dashed boxes in the left column (Scale bar, 10 µm) indicate the position of zoomed images shown in the other three columns. Scale bar, 10 µm. Plasma membrane staining by *Sp* Caveolin was observed in a small number of cells, but it was diffuse rather than punctate. (**L**) Colocalization of caveolin and Cavin1 was determined by Pearson’s Correlation analysis [*Hs* CAV1, n=25, *Sp* Caveolin, n=20, Sr Caveolin, n=25 regions of interest (ROIs) from at least two independent experiments]. One-way ANOVA with Tukey’s test (≥3 groups) was used to calculate p-values. n.s., not significant.

Previous cryo-ET studies of *h*-caveolae predate recent advances in cryo-EM ^24,64^. We, therefore, benchmarked *h*-caveolae formation using wild-type (WT) *Hs* CAV1-His_6_ expressed in *E. coli*. After 24 hours of induction, cells were vitrified, cryo-focused ion beam (cryo-FIB) milled, and tilt series collected on the milled lamellae (Table S1). Western blot confirmed stable expression of *Hs* CAV1-His_6_ (Figure 1D). In agreement with earlier results with MBP-CAV1 ^64^, cryo-ET revealed that *Hs* CAV1-His_6_, but not empty vector, induces inner membrane invaginations and *h*-caveolae formation (Figure 1E, F, and Video S1). On average, *E. coli* expressing *Hs* CAV1-His_6_ contained ∼38 *h*-caveolae per 0.01 µm³, with a mean diameter of ∼50 nm (Figure S2A-C). We also captured membrane budding events, suggesting the progression of *h*-caveolae formation (Figure S2D). Despite extensive tomogram analysis, the position of the *Hs* CAV1 complex in *h*-caveolae was difficult to visualize, suggesting the disc is deeply embedded in the membrane (Figure S1E).

We then performed a similar analysis using *Sp* or *Sr* Caveolins (Figures 1G, H, and Video S1). Despite *bona fide* expression (Figure 1D), the reconstructed tomograms showed no *h*-caveolae or inner membrane deformations for either variant. This lack of membrane remodeling was not due to defects in oligomerization or assembly, as we recently reported cryo-EM structures of the oligomeric (11-mer) *Sp* and *Sr* Caveolin complexes expressed in *E. coli* ^57^ (Figure 1B, C). Thus, while *Sp* and *Sr* Caveolins share the overall architecture of *Hs* CAV1, they do not induce membrane curvature in this model system.

### *Sp* and *Sr* Caveolins fail to induce caveolae in mammalian cells

We next tested whether *Sp* and *Sr* Caveolins can generate caveolae in CAV1^-/-^ mouse embryonic fibroblasts (MEFs), which lack caveolae. Myc-tagged *Hs*, *Sp*, or *Sr* Caveolins were transiently transfected and evaluated for colocalization with endogenous Cavin1. Caveolin and Cavin1 colocalization in cell surface puncta is indicative of caveolae biogenesis ^65,66^. *Hs* CAV1, as expected, colocalized strongly with Cavin1 at the plasma membrane, reflecting robust caveolae formation (Figure 1I, L). In most cells, *Sp* Caveolin staining was confined to spherical cytoplasmic structures positive for the lipid droplet marker Perilipin 3 and negative for Cavin1 (Figure 1J, L, and Figure S3A–F). Correspondingly, we observed an increase in lipid droplet size and number, as is typical of cells expressing lipid droplet-associated caveolins ^67,68^. *Sr* Caveolin trafficked to both the plasma membrane and intracellular vesicles and partially overlapped with Perilipin 3, but did not colocalize with Cavin1 (Figure 1K, L, and Figure S3C, D). Combined, these results show that, despite forming disc-shaped oligomers with central β-barrels, *Sp* and *Sr* Caveolins cannot recapitulate the ability of *Hs* CAV1 to induce *h*-caveolae formation or drive caveolae biogenesis.

### Residues at the rim of caveolin discs are key for caveolin-induced membrane remodeling

The unexpected inability of disc-shaped *Sp* and *Sr* Caveolin complexes to induce *h*-caveolae (Figure 1E–H) raises the question of which structural features are essential for membrane bending. Although the evolutionarily distant caveolin complexes share a conserved architecture, key differences, such as a charged ring on the hydrophobic face of *Hs* CAV1, disc curvature and diameter, and β-barrel length, may underlie their functional divergence (Figure 1A–C). To investigate these features, we used cryo-EM structures to engineer chimeras of *Sp* and *Sr* Caveolins containing domains from *Hs* CAV1. All mutants were expressed in *E. coli* and evaluated for *h*-caveolae formation by cryo-ET (Figure 2). Expression was confirmed by western blot, and structural integrity was assessed by negative-stain EM after detergent purification (Figure S4). We also tested each mutant’s ability to support caveolae biogenesis in CAV1^-/-^ MEF cells (Figure S5).

**Figure 2.**
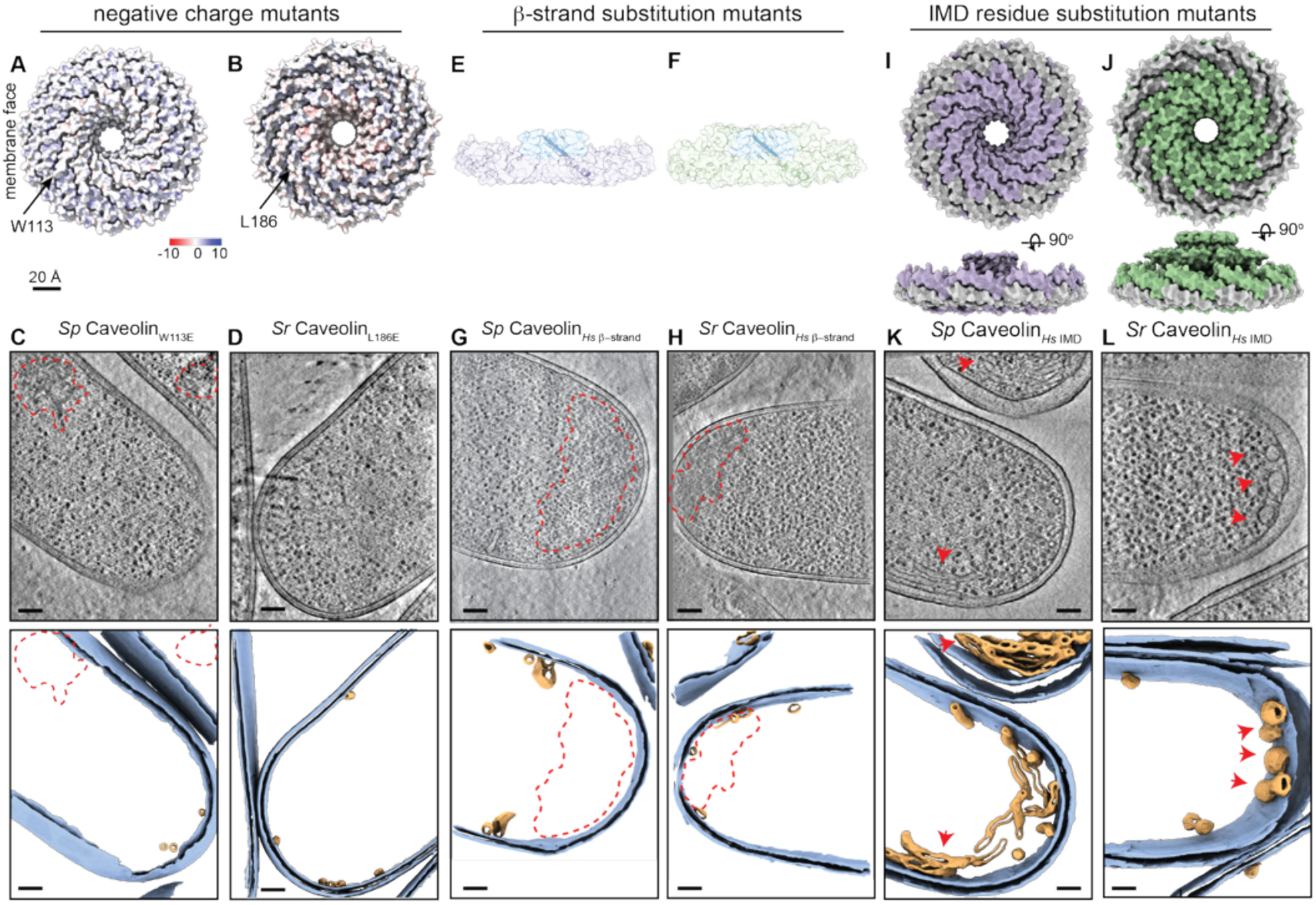
Substituting *Hs* IMD residues, but not other regions, in the *Sr* and *Sp* Caveolins alters their ability to sculpt membranes. (**A, B**) Space-filling model showing the charge distribution on the hydrophobic face of *Sp* and *Sr* Caveolin complexes. Arrows point to residue in equivalent position as *Hs* CAV1 Glu140. (**C, D**) Slices through representative deconvoluted tomograms (top panel) and corresponding segmentations (bottom panel) of *E. coli* expressing His_6_-tagged *Sp* Caveolin_W113E_ (C) or *Sr* Caveolin_L186E_ (D). (**E, F**) Cut through view of space-filling models of *Sp* Caveolin*_Hs_* _β-strand_ and *Sp* Caveolin*_Hs_* _β-strand_ mutant complexes with one β-strand shown as a ribbon diagram. (**G, H**) Slices through representative deconvoluted tomograms (top panel) and corresponding segmentations (bottom panel) of *E. coli* expressing His_6_-tagged *Sp* Caveolin*_Hs_* _β-strand_ (G) or *Sp* Caveolin*_Hs_* _β-strand_ (H). (**I, J**) Surface model of the predicted membrane facing surfaces of *Sp* Caveolin*_Hs_* _IMD_ and *Sr* Caveolin*_Hs_* _IMD_ mutant complexes rotated 90° around the x-axis. The equivalent position of the *Hs* CAV1 IMD residues in the *Sp* and *Sr* caveolins is shown in grey. (**K-L**) Slices through representative deconvoluted tomograms (top panel) and corresponding segmentations (bottom panel) of *E. coli* expressing His_6_-tagged *Sp* Caveolin*_Hs_* _IMD_ or *Sr* Caveolin*_Hs_* _IMD_. Red arrows mark the location of membrane invaginations. (C, D, G, H, K, and L) Bacterial membranes are shown in blue, *h*-caveolae or other small vesicles are shown in gold. Red dashed lines outline electron-dense regions that could represent inclusion bodies. Scale bars, 100 nm.

We first examined the role of the negatively charged ring of 11 glutamic acids (Glu140) on the hydrophobic surface of the *Hs* CAV1 disc, which is absent in *Sp* and *Sr* Caveolins ^57^. *Sp* and *Sr* Caveolin mutants with a similar ring of negative charges on the otherwise hydrophobic face of their discs (*Sp* Caveolin_W133E_ and *Sr* Caveolin_L186E_) did not induce *h*-caveolae or alter caveolin localization in CAV1^-/-^ MEFs (Figures 2A-D and S5B, E, G, J), with the caveat that detergent-purified *Sp* Caveolin_W113E_ failed to form stable complexes (Figure S4C). Conversely, neutralizing Glu140 in *Hs* CAV1 (*Hs* CAV1_E140A_) did not affect complex assembly (Figure S6A-E), *h*-caveolae formation in *E. coli* (Figure S7A, B), or co-localization with Cavin1 when expressed in CAV1^-/-^MEFs (Figure S7K, N). Therefore, the negatively charged ring on the membrane-facing surface of caveolin complexes is not required for caveolin-mediated membrane bending or caveolae formation.

Next, we examined the *Hs* CAV1 β-strand (residues 170–178), which forms the central β-barrel in the 8S complex. This barrel has a hydrophobic interior ^56^, is conserved across caveolins ^57^, and is important for *Hs* CAV1’s stability and function ^69–76^. While the diameter and the overall architecture of the β-barrels are generally conserved, the *Sp* and *Sr* β-strands are longer than and share low sequence conservation with the *Hs* CAV1 β-strand (0% and 14%, respectively) ^57^. To assess whether the *Hs* CAV1 β-strand length and/or sequence affects membrane sculpting, we replaced the *Sp* or *Sr* β-strands (residues 143–152 or 216–231, respectively) with the *Hs* CAV1 β-strand (residues 170–178) (Figure 2E, F). These substitutions did not affect complex stability (Figure S4F-J), did not recapitulate *h*-caveolae formation (Figure 2G, H), and did not increase colocalization with Cavin1 (Figure S5C, E, H, J). In the reciprocal experiment, replacing the *Hs* CAV1 β-strand with those from *Sp* or *Sr* Caveolins (Figure S7C, D) did not affect complex assembly (Figure S6F-J), *h*-caveolae formation in *E. coli* (Figure S6D, E and Figure S7E, F), or Cavin1 co-localization in CAV1^-/-^ MEFs (Figure S7L–N). These findings indicate that neither β-strand sequence nor length affects membrane bending or caveolae formation by *Hs*, *Sp*, or *Sr* Caveolins.

Finally, we assessed the role of IMD residues in the SR of *Hs* CAV1. Akin to the conserved β-barrel, the α-helical SR is a conserved structural feature among evolutionarily distant caveolins but shares little sequence similarity across the species examined here ^57^. IMD residues do not form a distinct domain, but rather are part of the helical SR located at the rim of the *Hs* CAV1 disc (Figure S1). To test the importance of the residues in the “rim” of the caveolin disc, we replaced residues corresponding to the IMD in the *Sp* and *Sr* Caveolins (residues 75-107 and 148-180, respectively) with the *Hs* CAV1 IMD residues (residues 102-134) (Figure 2I, J). Replacement of these IMD residues changes the ability of *Sp* Caveolin*_Hs_* _IMD_ and *Sr* Caveolin*_Hs_* _IMD_ to sculpt membranes (Figure 2K, L): expression of *Sp* Caveolin*_Hs_* _IMD_ induced tubular-shaped vesicles (red arrows, Figure 2K), while expression of *Sr* Caveolin*_Hs_* _IMD_ caused globular invaginations that remained attached or closely associated with the bacterial cytoplasmic membrane (red arrows, Figure 2L). While swapping *Hs* CAV1 IMD residues into *Sp* and *Sr* Caveolins changed their membrane-bending properties in *E. coli*, the subcellular distributions of these mutants in CAV1^-/-^MEFs were similar to WT *Sp* and *Sr* Caveolin (Figure S5D, E, I, J). The reciprocal *Hs* CAV1 mutants replacing the *Hs* IMD residues with those from *Sp* and *Sr* Caveolins (Figure S7G) exhibited reduced *Hs* CAV1 protein expression (Figure S6K), disrupted complex formation (Figure S6L-O), and a failure to produce *h*-caveolae in *E. coli* (Figure S7H-I). Since the *Hs* CAV1 IMD mutants were not structurally stable, we did not test their ability to drive caveolae formation in CAV1^-/-^ MEF cells. Overall, these results show that the residues near the rim of the caveolin discs contribute to caveolin-driven membrane sculpting; however, the substitution of the *Hs* IMD residues into the *Sp* and *Sr* Caveolins is not sufficient to drive the formation of characteristic spherical 50-100 nm *h*-caveolae induced by *Hs* CAV1 (Figure 1B).

### MD simulations identify a connection between the spatial hydrophobicity of the complex and membrane deformation

We previously showed that *Hs* CAV1 embeds in a single leaflet of the lipid bilayer and associates with highly curved membrane regions, irrespective of lipid composition ^58^. To test whether *Sp* and *Sr* Caveolins (PDB: 9DN0 and 9DN1, respectively) exhibit similar behavior, we performed coarse-grained MD simulations of the proteins embedded in a simple bilayer model composed of POPC (1-palmitoyl-2-oleoyl-sn-glycero-3-phosphocholine) and analyzed the effects on bilayer shape and protein conformation. Consistent with our previous observations ^58^, *Hs* CAV1 adopted a funnel-like conformation (Figure S8A, B), and was associated with regions of large positive curvature in the distal leaflet (Figure 3A and Figure S8C). In contrast to *Hs* CAV1, the *Sp* and *Sr* Caveolin complexes did not change conformation and associated with regions of the membrane that either remained flat or curved away from the complex (Figure 3B, C, and Figure S8B, C). These results are consistent with our experimental findings that, unlike the *Hs* CAV1 complex, *Sp* and *Sr* Caveolin oligomers do not bend membranes (Figure 1G, H).

**Figure 3.**
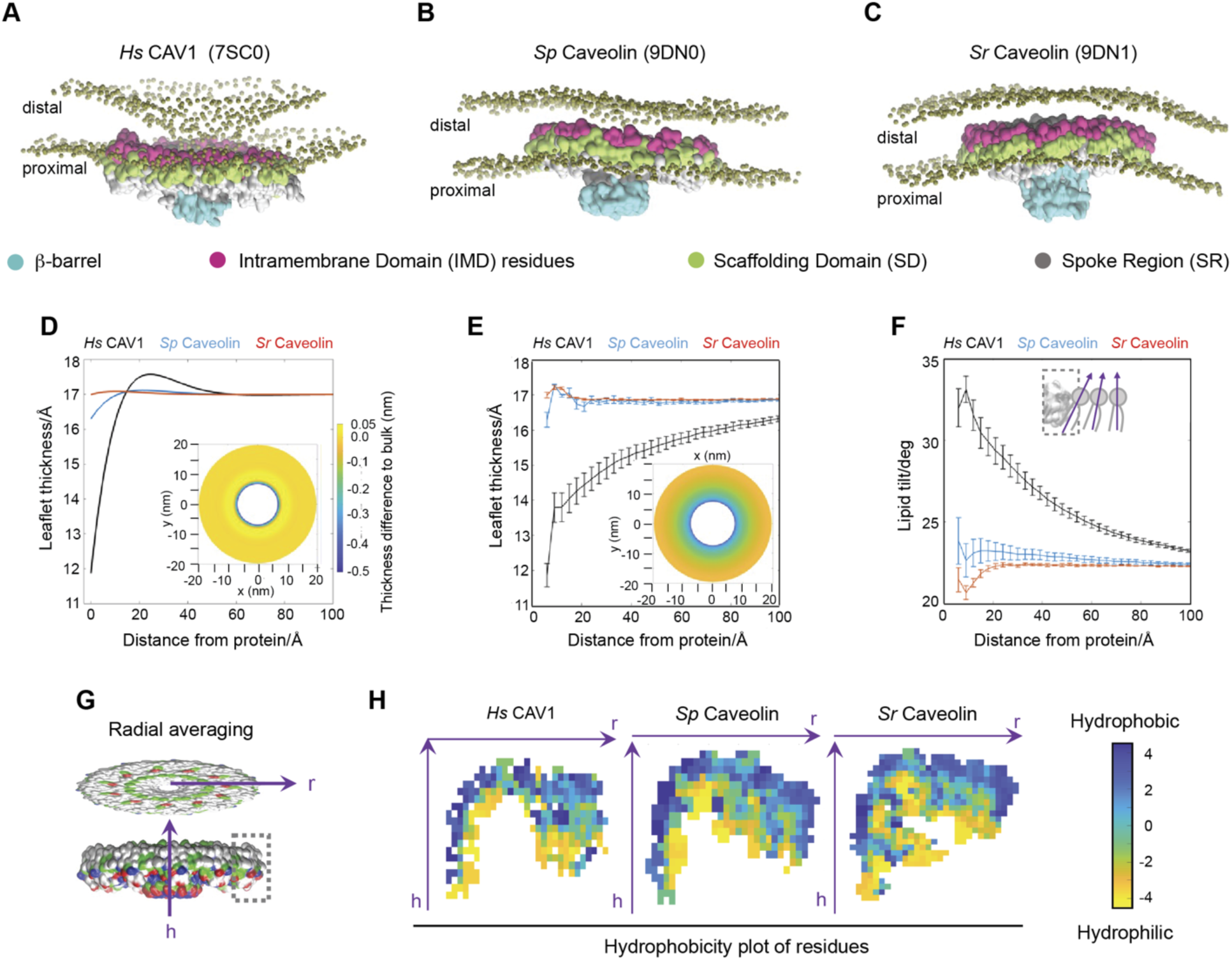
MD simulations recapitulate differences in the membrane-bending propensities of Caveolin complexes. (**A-C**) Simulation snapshots of *Hs* CAV1 (A), *Sp* Caveolin (B) and *Sr* Caveolin (C) complexes in a POPC membrane. Lipid phosphate groups within 50 Å of the caveolin complex are shown as tan spheres. Proximal and distal leaflets are labeled. The complexes are shown in surface representations. β-barrels shown in green, scaffolding domain (SD) shown in blue, intramembrane domain (IMD) residues shown in red, and shared residues between the SD and IMD residues shown in yellow. (**D-E**) Proximal leaflet thickness predicted from continuum theory (D) and measured from simulations (E) for the three complexes as a function of radial distance from the caveolin complex boundary. *Hs* CAV1 complex (7SC0), *Sp* Caveolin complex (9DN0), and *Sr* Caveolin complex (9DN1) shown in black, light blue, and red, respectively. Insets are 2D heatmaps showing the corresponding thickness changes relative to the bulk for the centered *Hs* CAV1 complex. Note that the theoretical model penalizes only thickness changes and does not consider lipid tilt. Color bar shown for panel in D is the same for panel in E. (**F**) Radially averaged tilt of proximal leaflet lipids as a function of distance from the three caveolin complexes. The inset shows the edge of the *Hs* CAV1 disc and lipid angles being measured (purple arrows). (G-H) Hydrophobicity mapping protocol (**G**) and resulting hydrophobicity maps (**H**) for the three caveolin complexes using the Kyte-Doolittle residue hydrophobicity scale. Note the sharper boundary between hydrophilic (yellow) and hydrophobic (blue) residues on the outer rim of *Hs* CAV1 complex compared to the *Sp* and *Sr* Caveolin complexes. Protein structure in (G) is colored by residue hydrophobicity: polar green, basic blue, acidic red, and nonpolar white; dashed box indicates the region of the protein-lipid boundary illustrated in the inset of (F); h, height; r, radius.

To understand the source of these differences, we examined the dynamics of the lipids around the protein. Since each caveolin complex displaced similar but slightly different numbers of lipids from one leaflet (see Methods), we first verified that the distinct morphological responses in the simulations were not due to the underlying interleaflet lipid imbalances (Figure S8E-J). We then analyzed the conformational dynamics of the lipids in the leaflet across from *Hs* CAV1 (the distal leaflet). A recent theoretical model proposed that unfavorable interactions between distal leaflet lipid tails and the hydrophobic surface of the complex cause the lipids to splay their chains, thereby contributing to curvature generation ^63^. However, while the curvature in the *Hs* CAV1 bilayer was different from that in the *Sp* and *Sr* Caveolin bilayers (with correlations of -0.2 and - 0.1, respectively), the distal lipid chain splay was similar (correlations of 0.6 and 0.4), and the correlation between curvature and splay across the three systems ranged from -0.4 to +0.6 (Figure S8K-N). These findings indicate that lipid splay alone is insufficient to explain the differences in membrane bending by caveolins.

In contrast, the lipids in the *proximal* leaflet—i.e., the leaflet in which the caveolin complex is embedded—showed a clear difference between the three variants. In particular, relative to the thickness of an unperturbed leaflet in a protein-free bilayer (17± 0.1 Å), the leaflet at the protein-lipid boundary was significantly thinner around *Hs* CAV1 (11.8 ± 0.3 Å) compared to *Sp* and *Sr* Caveolin (16.3 ± 0.2 and 17 ± 0.1 Å, respectively; Figure S9). Thus, *Hs* CAV1 complex, but not the *Sp* or *Sr* Caveolin complexes, substantially thins the surrounding leaflet at the protein-membrane boundary.

We wondered whether changes in membrane thickness at sites of caveolin complex insertion might explain some key features of membrane bending. To test this, we developed a simplified continuum model in which the elastic energy coupled to curvature generation was penalized by changes in membrane thickness (see Methods). We constrained the leaflet thicknesses at the protein-lipid boundary and in the bulk bilayer to those obtained from MD simulations and calculated the minimum-energy states associated with membrane bending through lipid thinning (Figure 3D). Our calculations revealed that changes in lipid thickness at the lipid-protein interface can indeed generate curvature within the continuum formulation. They also showed a predicted longer decay of lipid thickness to bulk in the *Hs* CAV1 system than in the *Sp* and *Sr* Caveolin trajectories (Figure 3D). However, for the latter two variants, these results provided a reasonable approximation of the actual thickness profiles obtained from the MD simulations (Figure 3E), whereas the predicted and actual deformations deviated significantly for *Hs* CAV1 (insets and black curves in Figure 3D, E). This suggests that while changes in lipid thickness contribute to membrane bending, additional factors must be present. Indeed, when we analyzed the dynamics of the lipids in greater detail, we found that the average tilt of proximal leaflet lipids relative to the three complexes differed substantially (Figure 3F). Notably, both lipid tilt and leaflet thickness were not fully converged even up to 100 Å away from the protein. This indicates that the overall perturbation of proximal leaflet lipids induced by a single complex can extend over long distances and likely plays a dominant role in determining the membrane’s deformation energy and, consequently, the extent of curvature.

To identify the molecular drivers of these distinct changes in lipid dynamics, we examined protein-lipid interactions at the caveolin-membrane boundary in detail (Figure S9). Analysis of the residues in direct contact with proximal lipid headgroups showed that they are located mainly in the SD and include both charged and non-charged residues in all caveolin variants (Figure S9G-I). For the *Sp* and *Sr* Caveolin complexes, these interactions resulted in the outer rim being mostly buried by surrounding lipids (Figure S9B-C), as revealed by their low solvent-accessible surface area (SASA; Figure S9E-F). In contrast, the lipid headgroups around *Hs* CAV1 are positioned closer to the bilayer interior on the outer rim of the complex (Figure S9A), resulting in greater solvent exposure for multiple residues along the disc periphery (Figure S9D). Importantly, structural analysis revealed that in the *Hs* CAV1 complex, non-hydrophobic residues are spread all along the rim, whereas in the *Sp* and *Sr* Caveolin complexes, they are localized towards the cytoplasm-facing part while the rest of the rim is mostly hydrophobic (Figure 3G-H). These distinct hydrophobicity profiles are fully consistent with the observed protein-lipid interactions and provide a possible explanation for why polar lipid headgroups preferentially interact with the parts of the rim of *Hs* CAV1 closer to the bilayer interior (Figure S9A-C).

### *h*-caveolae are not polyhedral

Several models have been proposed for how caveolins pack within caveolae and *h*-caveolae, including the possibility that caveolae membranes adopt a polyhedral geometry with *Hs* CAV1 complexes defining their flat surfaces ^27,56,62,64,77,78^. To test this idea, we attempted to localize *Hs* CAV1 complexes in isolated and vitrified *h*-caveolae using cryo-ET but were unable to confidently identify them because there was no discernible density outside the membranes, as expected for a largely membrane-embedded complex. To circumvent this limitation, we analyzed *h*-caveolae formed by *Hs* CAV1-mVenus, which we previously showed creates a distinctive “fan-like” density extending from the β-barrel ^69^ (Figure 4A, B). We first confirmed that *Hs* CAV1-mVenus expression in *E. coli* induced *h*-caveolae similar to *Hs* CAV1-His_6_. Tomograms showed that *E. coli* expressing *Hs* CAV1-mVenus, but not mVenus alone, caused membrane invaginations and *h*-caveolae formation (Figure 4C, D). Cells expressing *H*s CAV1-mVenus contained an average of ∼31 *h*-caveolae per 0.01 µm³ cell volume with a mean diameter of ∼37 nm (Figure 4E-G). We also captured snapshots of budding events at the inner membrane (Figure 4H). However, despite the presence of the mVenus tag, the position of these complexes within *h*-caveolae, *in situ,* was not discernible in the tomograms.

**Figure 4.**
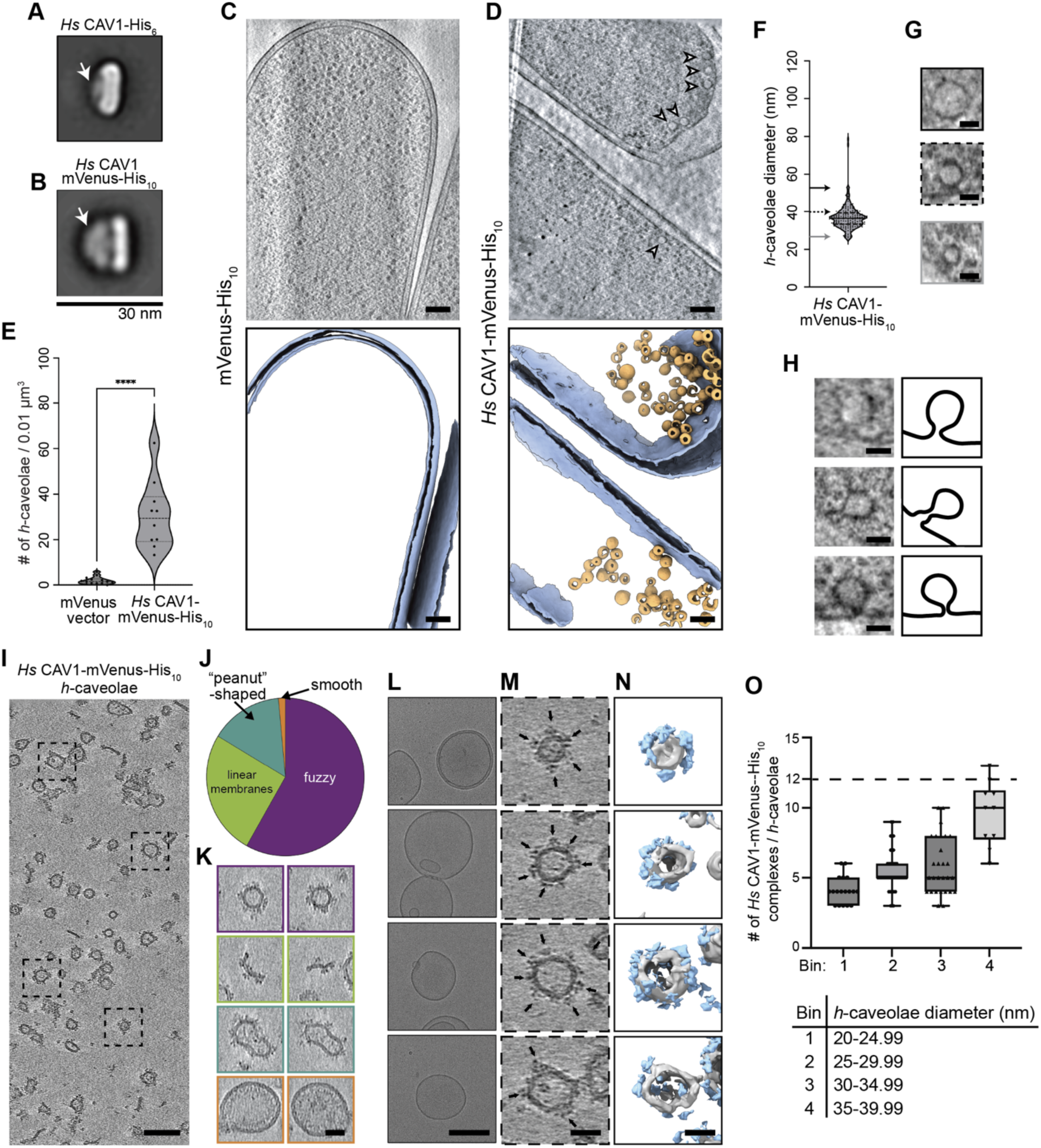
Cryo-ET analysis of *Hs* CAV1-mVenus reveals the number of complexes per *h*-caveolae and differences in morphology of cellular versus purified *h*-caveolae. (**A-B**) Representative examples of 2D class averages of negative stained detergent-solubilized *Hs* CAV1-His_6_ (A) and *Hs* CAV1-mVenus-His_10_ complexes (B). Scale bar, 30 nm. Arrow points to the C-terminus of the β-barrel. Averages modified from Han et al., Science Advances, DOI 10.1126/sciadv.abc6185 [2020], AAAS. (**C, D**) Representative slices through the tomogram and corresponding segmentation of *E. coli* expressing mVenus-His_10_ vector (C) or *Hs* CAV1-mVenus-His_10_ (D). Scale bars, 100 nm. White arrows mark *h*-caveolae invaginations at the inner membrane. In segmentations, bacterial membranes are shown in blue, and *h*-caveolae are shown in gold. (E) A violin plot showing the number of *h*-caveolae found per 0.01 µm^3^ of cellular volume in *E. coli* expressing the mVenus vector or *Hs* CAV1-mVenus-His_10_. **** *P* = <0.0001. Data are from 14 mVenus vector tomograms; 10 CAV1-mVenus-His_10_ tomograms. (**F**) A violin plot of the diameter of 170 *h*-caveolae visualized in 4 tomograms of *E. coli* expressing *Hs* CAV1-mVenus-His_10_. (**G**) Representative examples of *h*-caveolae imaged in the tomograms of *E. coli* expressing *Hs* CAV1-mVenus-His_10_. Outline around each image (black, dashed, and grey) corresponds to an arrow (black, dashed, and grey) showing its position in the violin plot shown in panel F. Images are the sum of 13 slices of the tomogram. Scale bar, 30 nm. (**H**) Representative image (right) and cartoon outline (left) of *h*-caveolae forming from the bacterial inner membrane. Images are a sum of 13 slices from the tomogram. Scale bars, 30 nm. (**I**) Slice through a deconvoluted tomogram of purified *h*-caveolae from *E. coli* expressing *Hs* CAV1-mVenus-His_10_. Four *h*-caveolae are highlighted by dashed boxes. Scale bar, 100 nm. (**J**) Pie chart of morphologies (%) observed in the tomograms of purified vesicles from *E. coli* expressing *Hs* CAV1-mVenus-His_10_. N= 349. (**K**) Examples of various morphologies quantified in panel J (linear membranes, peanut-shaped, fuzzy, and smooth). Images are a sum of 13 slices from the tomogram and outlined in the colors shown in panel J. Scale bars, 50 nm. (**L**) Representative cryo-EM images of vitrified liposomes composed of POPC. Scale bars, 150 nm. (**M**) Image of each *h*-caveolae boxed in panel G. Each image is a single slice of the tomogram. Black arrows point to a mVenus-His_10_ density protruding from the *h*-caveolae. (**N**) Corresponding segmentation of the *h*-caveolae shown in K. Membranes shown in grey and density for mVenus-His_10_ shown in light blue. Scale bars, 50 nm. (**O**) Number of *Hs* CAV1-mVenus-His_10_ complexes per *h*-caveolae of a particular diameter. Diameters included in each bin are listed. One *h*-caveolae was observed with a diameter greater than 40 nm (data not shown). N=380. Dashed line corresponds to 12 caveolin complexes/h-caveolae, the predicted number of complexes required for the polyhedral organization of caveolin complexes in a ∼50 nm *h*-caveolae.

We thus analyzed isolated and vitrified *h*-caveolae with *Hs* CAV1-mVenus by cryo-ET (Figure 4I–O). As anticipated, tomograms of *h*-caveolae from *Hs* CAV1-mVenus expressing cells displayed distinctive “fan-like” densities protruding from the membranes (Figure 4I). Notably, purified *h*-caveolae exhibited greater size and shape variability than *in situ*, where the *h*-caveolae were more uniform in size and mostly spherical in shape (Figure 4D, I–K). >50% of isolated and vitrified *h*-caveolae appeared as spherical vesicles with fuzzy densities surrounding their exterior (Figure 4J, K), with the remainder being a heterogeneous mix of sizes and morphologies (Figure 4D, I-K). As a control, vitrified protein-free liposomes were examined and found to be consistently spherical with smooth membranes (Figure 4L). Using a neural network-based algorithm, we mapped the locations of mVenus tags in “fuzzy” round *h*-caveolae (Figure 4M–O). We detected between 3-13 *Hs* CAV1-mVenus complexes in these *h*-caveolae (Figure 4O), although this may be a slight underestimation due to the missing wedge in cryo-ET data. To assess the relationship between *h*-caveolae size and the number of complexes, we grouped *h*-caveolae by diameter and found considerable variation in the number of *Hs* CAV1 complexes even among similarly sized *h*-caveolae. However, larger *h*-caveolae generally contained more *Hs* CAV1 complexes (Figure 4O). These complexes did not appear to adopt a uniform arrangement within the *h*-caveolae, which lacked a polyhedral geometry.

### The membrane-embedded *Hs* CAV1 complex is funnel-shaped and distorts the lipid bilayer

Since the *Sp* and *Sr* β-strands did not affect *Hs* CAV1 complex architecture or ability to form *h*-caveolae and caveolae (Figure S6D-J, Figure S7C-F, and Figure S6L-N), we decided to use the additional 10 Å height of the *Sr* β-barrel to localize and analyze the structure of *Hs* CAV1 complexes in membranes. To this end, we engineered an *Hs* CAV1 mutant with an extended β-barrel by adding *Sr* residues 225–233 to the C-terminus of Hs CAV1 (*Hs* CAV1_β-ext_). Control experiments confirmed that *Hs* CAV1_β-ext_ forms disc-shaped complexes with a longer β-barrel (Figure S10A-C), assembles *h*-caveolae in *E. coli* (Figure S10D-F), and co-localizes with Cavin1 in CAV1^-/-^ MEFs (Figure S10G-I).

We used single particle cryo-EM to determine a 4.1 Å resolution structure of the *Hs* CAV1_β-ext_ caveolin complex *in situ* in isolated *h*-caveolae. The resolution of the map allowed us to build a model encompassing the α-helical regions of the disc and rim along with the β-strand (residues 78-177); however, the density for the N-terminal region (residues 49-77) and eight additional *Sr* Caveolin residues appended to the *Hs* CAV1 β-barrel was not well resolved for *de novo* model building (Figure 5A-F, Figure S10J, Figure S11, and Table S3). The overall architecture of the membrane-embedded complex, composed of 11 spiraling α-helical protomers organized into a disc with a protruding β-barrel, is similar to *Hs* CAV1 in detergent micelles (Fig. 5A- F). However, we observed that the disc adopts a distinct funnel-shaped conformation in membranes *in situ* (Figure 5G-I), which distorts the trajectories of the distal and proximal leaflets of the lipid bilayer, as seen in the 2D averages (Figure 5G). We also observed strong density reminiscent of the distal leaflet that extends over the membrane-inserted side of the β-barrel (Figure 5G). Comparing the detergent-solubilized structure to the membrane-embedded structure shows that, in the context of the membrane, the α-helices that form the disc and rim (α1-5) adopt different conformation. While all five α-helices shift away from the distal leaflet towards the proximal leaflet, those closer to the β-strand show the largest changes (Figure 5I-K and Video S2). The combination of these individual shifts, together with their direction, leads to the disc adopting a funnel shape and positions the β-barrel further into the cytoplasm (Figure 5H, I).

**Figure 5.**
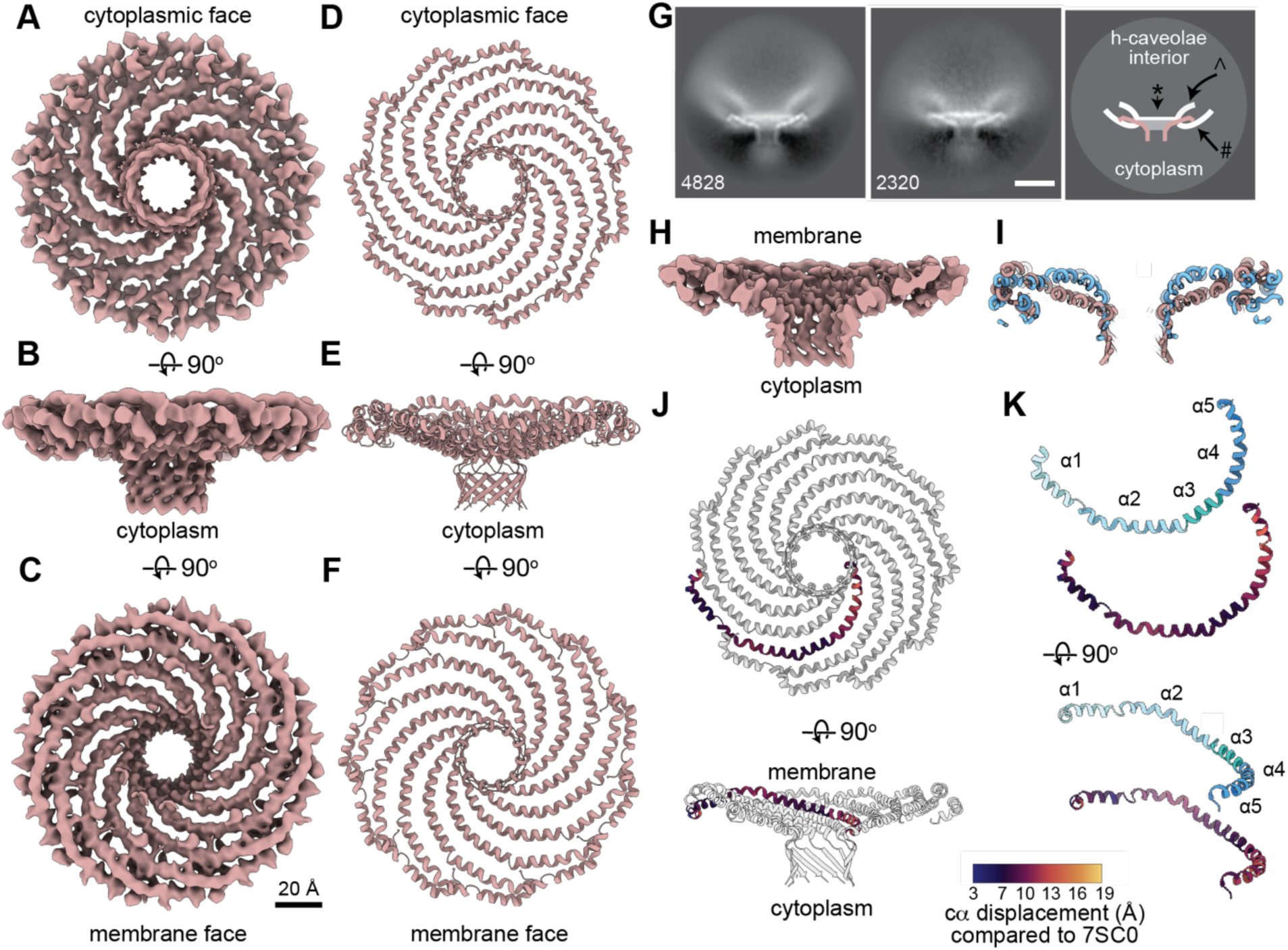
The *Hs* CAV1 complex adopts a funnel-shaped conformation in membranes. (**A-C**) 4.1 Å resolution 3D map of membrane-embedded *Hs* CAV1_β-ext_ complex with 11-fold symmetry applied, with three orthogonal views around the x-axis. Scale bar, 20 Å. (**D-F**) Model of the of membrane-embedded *Hs* CAV1_β-ext_ complex (residues 78-177) shown in the same orientations as panels A-C. (**G**) 2D class averages of *Hs* CAV1_β-ext_ in *h*-caveolae membranes (left and middle panels). Cartoon representation of 2D averages (right panel). *Hs* Cav1_β-ext_ complex shown in rose, membranes shown in white, and amorphous lipid density shown in transparent grey. ^, distal leaflet; #, proximal leaflet; *, density for lipids that extends across the membrane-inserted side of the β-barrel. Scale bar, 50 Å. (**H**) Slice through the map of *Hs* CAV1_β-ext_ complex in the same orientation as panel B, highlighting the densities of the complex (rose). (**I**) Comparison of slices of the models for detergent-solubilized *Hs* CAV1 complex (PDB 7SC0, blue) and membrane-embedded *Hs* CAV1_β-ext_ complex (rose), shown in the same orientation as panel H. (**J**) Orthogonal views of a model membrane-embedded *Hs* CAV1 _β-ext_ complex with one protomer colored by the relative displacement between Cα residues when comparing membrane-embedded and detergent-solubilized *Hs* CAV1 structure. Residues in the protomer are colored by Cα displacement in Å, with darker shades (purple) representing smaller changes and lighter shades (yellow) indicating larger changes. (**K**) Top: orthogonal views of a-helices 1-5 (residues 81-171) of a single protomer of the membrane-embedded *Hs* CAV1_β_-ext complex shown in panel J with a color scheme: α1 (residues 81–100), pale turquoise; α2 (residues 101-134), sky blue; a3 (residues 135-141), light sea green; α4 (residues 142-156), steel blue; and α5 (residues 157-168), dodger blue.Bottom: Relative displacement of the Cα residue between the membrane and detergent-solubilized models. Residues in the protomer are colored by Cα displacement in Å, with darker shades (purple) representing smaller changes and lighter shades (yellow) indicating larger changes.

## Discussion

Caveolins from evolutionarily distant species assemble into broadly similar disc-shaped oligomers, yet our findings demonstrate that this shared architecture alone does not drive membrane bending. While *Hs* CAV1, *Sp* Caveolin, and *Sr* Caveolin have low sequence similarity, all three assemble into amphipathic discs with central β-barrels. Despite this structural conservation, only *Hs* CAV1 forms *h*-caveolae in *E. coli* and caveolae in mammalian cells. Mutational and computational analyses revealed that the residues in the rim of the caveolin disc and their interactions with lipids determine the extent of deformation of the adjacent bilayer leaflet and consequently, membrane curvature. The cryo-EM structure of the *Hs* CAV1 complex in membranes shows the disc adopts a funnel-shaped conformation rather than the flat conformation found in detergent micelles. Thus, our results underscore the critical interplay between disc architecture, residue identity, and lipid interactions for caveolin-induced membrane sculpting.

### Properties of the Hs CAV1 complex required for membrane bending

Residues previously assigned to the SD and IMD have been linked to membrane curvature and caveolae formation ^24,66^; however, the underlying mechanism remains unclear, in part because the role of these residues in the *Hs* CAV1 structure is not well understood. From our cryo-EM analyses of caveolin complexes and structural predictions of evolutionarily diverse caveolins, we now know that each protomer contains five continuous α-helices (α1-α5). Helices α2 to α5 form the conserved SR that generates the flat surface of the disc ^56,57^ (Figure S1C). The outer rim of the *Hs* CAV1 complex disc is formed by helices α1 and α2, and includes residues previously designated as forming the SD (residues 82–100) and IMD (residues 101–134) (Figure S1A, C). According to our cryo-EM-base model, the IMD residues span structural elements contributing to both the SD and SR, rather than corresponding to a single structurally defined domain (Figure S1B, C). Here, we show that the location of these residues at the caveolin disc rim are critical for membrane bending.

Our analyses reveal that caveolin’s interactions with lipids in its surrounding proximal leaflet are essential for generating membrane curvature. In particular, the distribution of non-hydrophobic residues in the rim of the disc governs the positioning of these lipids, which may lead to varying degrees of local leaflet deformation. For example, in the experimentally determined caveolin structures, both lysine and arginine, in addition to other polar non-charged residues, are present in the rim of *Hs* CAV1 but not in *Sp* or *Sr* Caveolin. These residues promote favorable interactions of cytoplasmic lipid headgroups with the part of the complex closer to its membrane-embedded surface (Figure S1E and Figure S9A). Coupled with the relatively flexible α-helices of the *Hs* CAV1 complex that connect the disc to the β-barrel, the corresponding thinning of the leaflet facilitates the funnel-like conformational rearrangement of the complex (Figure S8A-B and Figure 5). Consequently, the extracellular leaflet curves towards the complex, as observed in both our MD simulations and cryo-EM studies (Figure 3A and Figure 5G). Indeed, structural predictions indicate that swapping the IMD regions of *Sp* and *Sr* Caveolins with those of *Hs* CAV1 results in both thinning of the ‘neck’ of the complex and a decrease in the overall hydrophobicity of the outer rim (Figure S12). This helps explain why the domain swaps impart curvature-inducing abilities to the non-human caveolin variants (Figure 2I-L).

Our results also disentangle the structural features of caveolin complexes redundant for curvature generation. For instance, the charged residues on the hydrophobic surfaces of the discs, residues forming the β-strand, or the length of the β-strand, do not affect caveolin’s ability to bend membranes or form caveolae. These findings are consistent with previous deletion analyses that identify regions critical for *h*-caveolae formation ^24^.

### Mechanisms of caveolin-induced membrane bending

Our data challenge several prevailing models of how caveolins bend membranes. First, the classic “wedging” model proposes that the IMD forms a structurally distinct α-helical hairpin structure that inserts into the lipid bilayer, inducing membrane curvature. Our recent structural studies revealed that these residues instead contribute to α-helices in the SR and SD, which together form the flat outer rim of the caveolin disc, deeply embedded in the cytoplasmic (proximal) membrane leaflet (Figure 5G) ^56^. Based on our current findings, we propose a “polar rim” model in which the hydrophobicity profile of these rim residues determines whether the complex can bend membranes, based on their interactions with lipids in the cytoplasmic leaflet. Notably, our structural model of the membrane-embedded *Hs* CAV1 complex reveals that the rim region adopts a distinct conformation *in situ,* in membranes, compared with detergent micelles *in vitro*, supporting the conclusion that this region of the complex interacts closely with lipids. Thus, while the IMD residues do not form a hairpin that inserts into the bilayer, their location at the rim of the *Hs* CAV1 disc makes them indispensable for membrane bending, providing a new molecular logic for the importance of these residues in caveolin complexes and their role in regulating membrane curvature.

A second, more recent, model proposes that *h*-caveolae and caveolae adopt a polyhedral geometry, with the flat, disc-shaped *Hs* CAV1 complexes forming the flat faces ^24,27,56,62,64^. This model has come under scrutiny because MD simulations indicate that the CAV1 complex prefers a funnel-shaped, rather than flat, conformation when interacting with membranes ^58,80^. In this study, we present multiple lines of experimental evidence that also challenge the polyhedral packing model. *h*-caveolae *in situ* are generally round (Figure 4D), with flat surfaces rarely observed; appearing primarily in isolated *h*-caveolae (Figure 4I-N). Additionally, the number of complexes and their localization in *h*-caveolae do not strictly correlate with *h*-caveola size (Figure 4O), suggesting that these complexes do not pack in a highly regular or geometric manner. Finally, we demonstrate that the *Hs* CAV1 complex indeed adopts a funnel shape when embedded within *h*-caveolae membranes Figure 5). Taken together, these findings indicate that, at least in *h*-caveolae, *Hs* CAV1 complexes are not organized in a regular polyhedral geometry. However, such a geometry may be imposed by other components of *bona fide* caveolae, such as the cavins ^77^.

A newer theoretical model postulates that the presence of an amphipathic disc in the inner leaflet of the membrane causes thinning of the extracellular leaflet, driven by the angle between the two lipid acyl chains – ‘lipid splay’ ^63^. Increased lipid splay is hypothesized to arise from unfavorable (relative to lipid-lipid interactions) lipid-protein interactions at the bilayer midplane and thus to contribute to membrane bending. The 2D class averages of the CAV1 complex embedded in *h*-caveolae show that the membrane thins where the distal leaflet overlays the disc and the proximal leaflet interacts with the cytoplasmic side of the disc (Figure 5G). This is also supported by recent coarse grain simulations of *Hs* CAV1 complexes in membranes of various lipid compositions ^58^. Our current MD simulations show that phospholipids in the extracellular leaflet, which make direct contact with the membrane-facing surface of the *Hs* CAV1, *Sr*, and *Sp* Caveolin complexes, can splay out their chains to a similar extent irrespective of the identity of the complex (Figure S8). Considering that only *Hs* CAV1 was observed to induce curvature both in our experiments and simulations, we conclude that while lipid splay of the extracellular leaflet may partially contribute to membrane bending, it is not the primary mechanism.

Our finding that *Hs* CAV1 can adopt two distinct conformations raises the possibility that the complexes may interconvert between a flattened and funnel shape, allowing them to reside in both flat and curved membrane environments ^81^. Such dynamics could potentially facilitate their ability to respond to triggers such as mechanical stress ^55,82^. How these events or the process of membrane bending may be assisted by collective interactions between caveolin complexes, including the formation of scaffolds and higher-order 70S complexes, remains to be uncovered^28,78,83,84^.

### Evolutionary and functional implications of caveolin diversity

While mammalian caveolins are well known for their central role in caveolae biogenesis, caveolins also have functions independent of caveolae ^85–87^. The inability of *Sp* and *Sr* Caveolins to bend membranes or form caveolae in mammalian cells may reflect a specific evolutionary adaptation of mammalian caveolins to promote caveolae biogenesis rather than a loss of conserved functionality. Caveolin’s ability to form a protein interaction scaffold on membranes, and a conserved central β-barrel that spans the proximal leaflet of the bilayer, may represent the structural features required to support ancient caveolin functions, such as facilitating signaling pathways and/or supporting lipid homeostasis. Indeed, MD simulations suggest that cholesterol and other lipids are sequestered within the β-barrel domain, and mutations in this region in human caveolins are linked to lipid disorders and altered lipid droplet dynamics ^58,59,69–71,73,88^. Thus, the most evolutionarily conserved function of caveolins may predate their established role in caveolae biogenesis.

In summary, our work establishes that membrane curvature generation by caveolins requires not only oligomerization and amphipathic disc formation but also precise residue patterning at the disc rim and conformational flexibility in the context of the membrane. Our polar rim model highlights the importance of proximal leaflet deformation, thus identifying a new mechanism by which proteins remodel membranes. These insights advance our understanding of the molecular basis of caveolae biogenesis, metastability, and open new directions for exploring caveolin function in membrane remodeling and lipid regulation.

## Resource availability

### Lead contact

Further information and requests for resources and reagents should be directed to and will be fulfilled by the lead contact, Melanie D. Ohi.

### Data and Materials Availability

All reagents generated in this study are available from the lead contact upon completion of a materials transfer agreement. The map and model have been deposited in EMDB (EMD-XXXXX) and PDB (PDB-XXXX). All other data needed to evaluate the paper’s conclusions are included in the paper or the supplementary materials.

### Funding Sources

Research reported in this publication was supported by: NIH 1R01HL168258 (AKK), NIH 1R01GM151635 (AKK), NIH DP2GM150019 (SM), NIH NIGMS NIH R35GM158446 (PR), NIH S10OD030275 (MDO), NIH T32GM140223 (TSB), NIH T32GM007315 (SMC and SR), AHA 905705 (SMC), the Arnold and Mabel Beckmann Foundation imaging grant to the University of Michigan Cryo-EM Facility (MDO). SciLifeLab and Wallenberg Data Driven Life Science Program KAW 2024.0159 (MD), and Swedish Research Council Starting Grant VR 2025-04114 (MD). The content is solely the responsibility of the authors and does not necessarily represent the official views of the National Institutes of Health.

## Acknowledgements

We thank Yelena Peskova for expert technical assistance; Tim Baker and Wil Salmen for measuring differences between structures; and Brigitte Meyers for assistance at early stages of this study. We thank the U-M Cryo-EM facility staff for scientific and technical assistance. We thank Drs. Ilya Levental and Herman Fung for comments on the manuscript, and members of the Ohi and Kenworthy labs for helpful discussions.

