## Supplemental Data for "Structural basis of caveolin-driven membrane bending" (from another version of this preprint: version 1, 2026-02-08)

**for**

**This PDF contains:**

Materials and Methods

Supplementary Note 1

Figures S1-12

Tables S1-3

### Materials and Methods

#### Construct preparation

For *E. coli* expression, caveolin point mutants were generated using site-directed PCR mutagenesis as described previously [1] using pET20 WT *H. sapiens* CAV1, pET20 WT *S. purpuratus* Caveolin, or pET20 WT *S. rosetta* Caveolin as a template [2,3]. The primers used (Integrated DNA Technologies, IA) were as follows: *H. sapiens* CAV1 E140A CATGAAACTGTACTACGTGGAGCTGCGGGTGTTCAACCTG; *S. purpuratus* Caveolin W113E CATGAAACTGTACTACGTGGAGCTGCGGGTGTTCAACCTG; *S. rosetta* Caveolin L186E CAACCAGCTGATGTTCCTGGAGATCGCCAGCCTGGCCCC. Caveolin fusion proteins (IMD or β-barrel swaps) cDNAs were codon-optimized for expression in *E. coli* and were synthesized by Twist Biosciences, CA. These cDNAs were subcloned into pET20b (+) vector (Novagen, #69739) using NdeI and XhoI (New England Biolabs) restriction sites. All clones were confirmed by Sanger sequencing.

For mammalian expression, genes encoding the entire Caveolin polypeptide of *S. purpuratus* (UniProt Accession A0A7M7T4C2) and *S. rosetta* (UniProt Accession F2U793) were synthesized by Genscript, NJ and further cloned into the pcDNA3.1-myc plasmid using KpnI and SacII (New England Biolabs, MA) restriction sites. *Hs* CAV1-myc as described previously [4]. The IMD swaps were generated using the corresponding plasmids for the bacterial expression described above. Sequences encoding the *Sr* and *Sp* Caveolin human IMD were cloned into the pcDNA3.1-myc plasmid using the same restriction sites and primers. Primers are provided in Supplementary Table 2. All point mutations, as well as β-barrel swaps and an extension of *Hs* CAV1 with the portion of the *Sr* β-barrel, were introduced by the mutagenesis using the Quick Change Lightning kit (Agilent, #210518). The resulting plasmids were verified by Sanger sequencing (Genewiz/Azenta Life Sciences, NJ).

#### Western blotting

Before induction (T0) and after induction (T24), 1 OD of cells was pelleted at 5000× g, supernatant removed, and pellets resuspended in 2× SDS Laemmli sample buffer (Bio-Rad). Proteins were resolved by SDS-polyacrylamide gel and transferred to nitrocellulose membranes (Bio-Rad). Membranes were probed with monoclonal anti-6x His tag antibody (Sigma SAB2702218-) to detect His-tagged proteins and anti-*Bacillus subtilis* ICDH antibody (Kerafast ETU002) as a loading control, and membranes were visualized using horseradish peroxidase-conjugated secondary antibodies (Thermo Fisher Scientific) and SuperSignal™ West Pico PLUS Chemiluminescent Substrate (Thermo Fisher Scientific). The blots were imaged on a ChemiDoc™ Touch Imaging System (Bio-Rad).

#### Mammalian cell culture

Caveolin-1 knock-out (KO) MEFs (CAV1$^{-/-}$ MEFs) obtained from ATCC were cultured in Dulbecco's modified Eagle's medium (Invitrogen, #11995-065) (containing 10% fetal bovine serum (Invitrogen, #A5670701), 1% Pen/Strep at 37 °C, and 5% $CO_2$ in a tissue culture incubator.

#### Immunofluorescence Assays

CAV1$^{-/-}$ MEFs were seeded in 6 well plate containing 12 mm diameter, #1.5 thickness glass cover slips (Electron Microscopy Science, #72290-04). 5 hours post-seeding, cells were transfected with Lipofectamine 2000 (Thermo Fischer # 11668-027) with 2 µg of each of the indicated constructs. 20-24 h post-transfection, the cells were washed and fixed in 4% paraformaldehyde for 20 min. Cells were then washed three times in 100 mM glycine in PBS to quench the crosslinker. Samples were blocked and permeabilized with blocking buffer (1% BSA containing

0.1% saponin in 1× PBS) for 1 h at room temperature (RT) in a humidified chamber. Cells were co-stained with mouse anti-Myc (Cell Signaling Technology, #911B, 1:100 dilution in blocking buffer) and rabbit anti-cavin1/PTRF (Proteintech, #18892-1-AP, 1:100 dilution in blocking buffer) or with guinea pig anti-perilipin (Progen, #GP432) antibody for 2-3 h at RT in a humid chamber. After washing 3× in PBS, the cells were incubated in Alexa 488 and Alexa 546-conjugated secondary antibodies (488 goat anti-mouse IgG, Life Technologies, #A11029; 546 goat anti-rabbit IgG, Life Technologies, #A11035; 546 goat anti-guinea pig, Life Technologies, # A11074 ) at a dilution of 1:300 for 1 h at RT in a humid chamber. Coverslips were then washed 4-5 times in PBS and mounted on glass slides (Thermo Scientific, #3050-002) using ProLong Gold Antifade reagent with DAPI (Cell Signaling Technology, #8961S). Cells were imaged using a Zeiss LSM 880 confocal microscope with a Plan-Apochromat 63×/1.4 oil DIC M27 objective at RT using a confocal slice of 1 Airy unit. Signals from the green (Alexa 488) and red (Alexa 546) channels were collected with a 34-channel GAsP spectral detector. All images were collected at 16-bit depth. Contrasts were adjusted linearly with ImageJ and exported as TIFFs.

Colocalization analysis was performed in Fiji 2.16.0/1.54p; Java 1.8.0_172 [64bit] using the colocalization plugin. Briefly, images used for colocalization analysis were acquired at a 4× zoom factor. For each set of comparisons, ≥ 20 images from at least two independent experiments were used for analysis. Pearson correlation coefficients are reported in a box-and-whisker plot. To quantify lipid droplets, the number per cell under each condition was manually counted using the counter plugin in Fiji. The number of lipid droplets per cell was calculated by outlining individual lipid droplets and measuring their areas using the measurement function in Fiji. A one-way ANOVA with Tukey's test (≥3 groups) or an unpaired Student's *t* test (2 groups) was used to calculate *p*-values and plotted using GraphPad Prism 10.6.1 (892).

### Expression and purification of *h*-caveolae and caveolin complexes

Caveolins were expressed in *E. coli* BL21-DE3 using a described auto-induction method [3,5]. Briefly, monoclonal bacteria were cultured in MDG media for 20 hours at 37 °C and 250 rpm. Starter cultures were expanded in auto-inducing ZYM-5052 medium for 24 h at 25 °C and 300 rpm. After 24 h of induction, cells were collected for cryo-ET analysis. Purification of caveolin complexes was performed as described previously [3,6,7]. *E. coli* BL21-DE3 cells that were autoinduced for 24 h were pelleted and resuspended in a buffer of 200 mM NaCl, 20 mM Tris (pH 8.0). Following the addition of 1 mM Dithiothreitol (DTT) and phenylmethylsulfonyl fluoride (PMSF), cells were homogenized with a pressure homogenizer (Avestin). Cellular debris was pelleted and removed by centrifugation at 4 °C for 15 min at 14,000 *g*. The remaining sample was centrifuged for 1 h at 4 °C and 114,500-160,000 *g* to isolate the membranes. The isolated membranes were resuspended in a buffer containing 200 mM NaCl, 20 mM Tris (pH 8.0), and 1 mM DTT, then homogenized with a Dounce homogenizer. To solubilize the membrane, n-dodecyl- β-D-maltoside (DDM) (Anatrace) was added to a final concentration of 2%, and the membrane fraction was incubated while rocking for 2 h at 4 °C. Insoluble material was removed by centrifugation at 42,000 rpm for 35 min. The supernatant was added to a nickel-charged Sepharose column (Chelating Sepharose Fast Flow, GE Healthcare) that had been pre-equilibrated with a buffer of 200 mM NaCl, 20 mM Tris-HCl (pH 8.0), 0.05% DDM, 10 mM imidazole, and 1 mM DTT. The supernatant and equilibrated resin were incubated together with gentle agitation at 4°C for 2h. The resin was washed with a buffer of 200 mM NaCl, 20 mM Tris-HCl (pH 8.0), 0.05 DDM, 30 mM imidazole, and 1 mM DTT. The protein was eluted with a buffer of 200 mM NaCl, 20 mM Tris-HCl (pH 8.0), 0.05% DDM, 300 mM imidazole, and 1 mM DTT and further purified using a Superose 6 Increase 10/300 GL column (GE Healthcare) that was equilibrated with a buffer of 200 mM NaCl, 20 mM Tris-HCl (pH 8.0), 1 mM DTT, and 0.05% DDM. Fractions containing caveolins were examined by negative stain EM.

To isolate *h*-caveolae, the homogenized total membrane (before the solubilization step) was run over a nickel-charged Sepharose column pre-equilibrated with a buffer of 200 mM NaCl, 20 mM Tris-HCl (pH 8.0), 10 mM imidazole, and 1 mM DTT. The resin was washed with buffers containing increasing imidazole concentrations before elution with elution buffer (200 mM NaCl, 20 mM Tris-HCl, pH 8.0, 400 mM imidazole, 1 mM DTT). Elutions were dialyzed overnight in buffer containing 200 mM NaCl, 20 mM Tris-HCl (pH 8.0), and 1 mM DTT. Dialyzed elutions were used for vitrification.

Caveolin samples were prepared for negative stain EM using established methods [8]. Briefly, a PELCO easiGlow™ glow discharge unit (Fresno, CA, USA) was used to glow-discharge 200-mesh copper grids covered with carbon-coated collodion film (EMS, Hatfield, PA, USA) for 30 s at 10 mA. Caveolin samples (3.5 µL) were adsorbed to the grids and incubated for 1 minute at room temperature. Samples were washed with two drops of water, then stained with two drops of 0.7% (w/v) uranyl formate (EMS, Hatfield, PA, USA), and blotted until dry. A Morgagni transmission electron microscope (TEM) operated at 100 kV (Thermo Fisher Scientific, Waltham, MA, USA) was used to image the samples at a nominal magnification of 22,000x (2.1 Å/pixel).

Negative stain EM datasets were collected using a Tecnai Spirit T12 TEM operated at 120 kV (Thermo Fisher Scientific, Waltham, MA, USA). Images were collected at a nominal magnification of 30,000× (2.086 Å/pixel). Data was collected using SerialEM v4.0.8 on a 4 k x 4 k Rio camera (Gatan, Pleasanton, CA) at a defocus of -2.2 µm [9]. All image processing was performed in RELION 4.0.0 [10]. Approximately 1,000 particles were manually selected, 2D classified, and clear resulting classes were selected and used as references for particle picking on all micrographs. Particles were extracted with a box size of 128 × 128 pixels (267 Å × 267 Å). The extracted particles were classified into 20 classes. The *Hs* $CAV1_{E140A}$, *Sr* $Caveolin_{L186E}$, *Hs* $CAV1_{\beta\text{-ext}}$, *Sp* $Caveolin_{Hs\ IMD}$, and *Sr* $Caveolin_{Hs\ IMD}$ datasets had 132,996, 28,889, 50,882, 1,077, and 12,142 particles, respectively.

### Sample preparation of cryo-EM/ET studies

#### *E. coli for in situ cryo-ET*

Following 24 h induction of the *E. coli* expressing caveolins, 10 mL of bacterial cells (relative $OD_{600}$ of 1.2-1.9) were spun down at 500× g for 5 minutes, supernatant was removed, and bacterial pellets were resuspended in ZYM-5052 media to a relative $OD_{600}$ of 20. Copper grids, 200 mesh, with holey R 1/4 carbon film (Quantifoil Micro Tools GmbH) were glow-discharged for 30 seconds at 5 mA using an EasiGlow system (Pelco) 1 h prior to sample application. Grids were loaded into the environmental chamber of a Vitrobot Mark IV (ThermoFisher Scientific) set to 25 °C and 95% humidity. 3 µL of resuspended bacteria were pipetted onto the carbon surface of the grid, and excess moisture was immediately blotted away for 5 seconds at a blot force of 10. Whatman filter paper no. 1 was used to back-blot the grid, while a Teflon ring was placed on the sample-facing blot arm of the Vitrobot to simulate single-side blotting. Following blotting, grids were immediately vitrified by plunge-freezing in liquid ethane.

#### *Sample preparation for isolated h-caveolae*

Quantifoil R 2/2, 200-mesh Cu grids with an ultrathin carbon layer (Electron Microscopy Services) were glow-discharged for 30 s at 5 mA using an EasiGlow system (Pelco) before vitrification. 3.5 µL of purified and undiluted *h*-caveolae formed by either *Hs* CAV1-mVenus-$His_{10}$ or *Hs* $CAV1_{\beta\text{-ext}}$ was applied to a glow-discharged grid. The sample was incubated on the grid for 30 s, blotted with Whatman filter paper no. 1 for 5.5 s at a blot force of 15, and then plunge-frozen in a slurry of liquid ethane using a Vitrobot Mark IV (Thermo Fisher Scientific). The Vitrobot chamber was set to 4°C and 100% humidity.

### Cryo-focused ion beam milling

Vitreous frozen grids containing bacterial samples were clipped into cryo-FIB AutoGrids (Thermo Fisher Scientific). Clipped grids were loaded onto the stage of an Aquilos 2 FIB-scanning electron microscope (SEM) (Thermo Fisher Scientific) and maintained at ~-185 °C. Grids were sputter-coated with an inorganic platinum layer, then organometallic platinum was deposited using a gas injection system for 30 s, and finally sputter-coated with a second inorganic platinum layer to form a protective/conductive surface for milling. Initial cuts with the FIB to make tension-relief trenches and 5 μm lamellae were milled with a beam current of 1 nA at a milling angle of 6° [11]. Subsequent cuts to make rough (1 μm) lamella were done with beam currents of 100-500 pA. Milling progress and lamella integrity were assessed by SEM. Upon completion of all rough lamella milling, each lamella was revisited and polished to a fine lamella thickness of ~200 nm with the FIB at a beam current of 50 pA [12].

### Tilt series acquisition

Tilt series on the milled lamella were collected on a Titan Krios G4i (ThermoFisher Scientific) operating at 300 kV and equipped with a K3 direct-electron detector (Gatan Inc.) A BioQuantum energy filter with a slit width of 20 eV (Gatan) was used. Images were collected at a nominal magnification of 42,000×, corresponding to a pixel size of 2.122 Å. Tilt series were collected using SerialEM v4.0.3 with a modified dose-symmetric tilt scheme ranging from -60 to +60 degrees in 3° increments [13,14]. The total target accumulated dose was ~120 $e^-/Å^2$ per tilt series. 12 frames were collected per tilt, and the frames were motion-corrected and aligned on the fly using SerialEM [9].

### Tilt series alignment and tomogram deconvolution

Tilt series were aligned using with patch tracking in IMOD v4.11 or AreTomo v1.3.4 [13,15]. Bad tilts were removed using default dark image thresholding (value of 0.7) in AreTomo or manually in IMOD. Tilt series aligned in IMOD were reconstructed using weighted back-projection with a SIRT-like filter (15 iterations). Subsequently, tomograms were deconvoluted using custom MATLAB 2020b scripts as described previously [16]. All tilt-series datasets acquired are summarized in Supplementary Table 1.

### Tomogram quantification

The number of *h*-caveolae per cell was counted manually in IMOD v 4.11 [13], and vesicle diameters were determined by averaging three manual measurements of the vesicle's central slice. To determine cell volumes, *E. coli* cells were divided into the end of the cell, modeled by a sphere, and the body of the cell, modeled by a cylinder. Cell volumes were calculated by the following:

$$V_{cell} = C_V + D_V \ ,$$

Where $C_V$ is the volume of a cylinder, $C_V = \pi r^2 h$ and $D_V$ is the volume of a dome (half sphere) $D_V = \frac{2}{3}\pi r^3$. The thickness of each cell was measured manually three times along the Z-dimension using IMOD v 4.11[13], and the average of these measurements was taken as the cell diameter. The height of the cylinder portion of the cell was also measured in IMOD v 4.11 [13].

### Cryo-EM data collection for *Hs* CAV1 $_{\beta\text{-ext}}$ *h*-caveolae

Micrographs of the purified *Hs* CAV1 $_{\beta\text{-ext}}$ *h*-caveolae were acquired on a Titan Krios electron microscope (ThermoFisher Scientific) operated at 300 kV and equipped with a K3 Summit direct electron detector (Gatan) (Supplementary Table 3). A BioQuantum energy filter with a slit width of 20 eV (Gatan) was used. Images were collected at a nominal magnification of 105,000× (0.83 Å/pixel). Two datasets of the *Hs* CAV1 $_{\beta\text{-ext}}$ *h*-caveolae sample were collected, containing 8,150 and 6,244 micrographs using, SerialEM v4.0.3 [9]. Micrographs were collected with a total dose of 50 $e^-/Å^2$and a defocus range of -0.5 µm to -3 µm.

#### Image processing of *Hs* CAV1 $_{\beta\text{-ext}}$ *h*-caveolae

All image processing, including 2D classification and 3D refinements, was carried out in cryoSPARC v4.3.1 [17]. Briefly, two datasets of 8,150 and 6,244 micrographs (14,394 micrographs total) were acquired, and the micrographs were patch-motion corrected and patch CTF estimated. Following manual selection of ~600 particles, a Topaz picking model was trained on a subset of 2,045 micrographs and was then used to select particles in both datasets [18]. Following micrograph curation and the combination of datasets, 279,236 particles were extracted with a box size of 256 $pixels^2$ (214 Å x 214 Å), Fourier cropped to a box size of 100 $pixels^2$ (83 Å x 83 Å), and 2D classified into 100 classes. The 2 classes that clearly showed the *Hs* CAV1 $_{\beta\text{-ext}}$ complex in *h*-caveolae membrane were selected for further processing. These 2 classes contained 17,483 particles. Particles were re-extracted from 7,629 micrographs at a 350 $pixels^2$ (292 Å x 292Å) box size, resulting in 16,788 particles. These particles were 2D classified into 50 classes. Four classes corresponding to 10,853 particles were selected as inputs for a single-class *ab initio* 3D reconstruction with $C_{11}$ symmetry imposed. Using this *ab initio* reconstruction and the corresponding particles, a $C_{11}$ non-uniform refinement resulted in a 3D reconstruction of 4.39 Å. The particles from this refinement were symmetry expanded using $C_{11}$ point group symmetry. The symmetry expanded particles were input into a $C_1$ local refinement job with a loose mask over the complex but not the membrane density. This resulted in a map with a reported resolution of 4.05 Å (Supplementary Table 3). This map was sharpened with a *B* factor of -130 and used for model fitting.

The *Hs* CAV1 model determined in a detergent micelle (PDB: 7SC0) was first rigid-body fitted into the *Hs* CAV1 $_{\beta\text{-ext}}$ density in ChimeraX (v1.10.1) [19]. ISOLDE [20] was then used to relax the *Hs* CAV1 model into the *Hs* CAV1 $_{\beta\text{-ext}}$ density. The model was then further fit with ISOLDE and Coot [20,21] and it was iteratively refined in Phenix[22]. A lack of defined density at the N-terminal region of the map led to the removal of residues 49-77 from the *Hs* CAV1 $_{\beta\text{-ext}}$ model, resulting in the final *Hs* CAV1 $_{\beta\text{-ext}}$ model encompassing residues 78-177.

#### Molecular Dynamics Simulations

*Systems setup.* The complex structures of *Hs* CAV1 (PDB: 7SC0), *S. purpuratus* (PDB: 9DN0), and *S. rosetta* (PDB: 9DN1) Caveolins were taken from experimental Cryo-EM data [2,6]. Each protomer of the complexes contained residues 49 to 177 for 7SC0, residues 29 to 152 for 9DN0, and residues 79 to 231 for 9DN1. All protomers had charged terminal end caps.

Each complex was embedded in a POPC bilayer in coarse-grained representation. All systems were built using CHARMM-GUI [23-25]. The initial placement of each complex in the bilayer was determined with OPRLM [26] , while the number of lipids displaced from the proximal leaflet (in which the complex was embedded) was calculated by CHARMM-GUI. The final systems contained 2965 POPC lipids in the distal leaflet (across from the complex), and 2789, 2781, or 2787 POPC lipids in the proximal leaflet around the 7SC0, 9DN0, and 9DN1 complexes, respectively. Each protein-bilayer system was solvated with a 75 Å-thick water layer above and below the bilayer and with 0.15 M NaCl. In addition, we built a lipid-only bilayer (without protein) with either equal numbers of lipids in both leaflets (balanced POPC) or with the same number of lipids removed from one leaflet as in the corresponding complex-bilayer simulation (imbalanced POPC). The bilayer-only systems were solvated with 0.15 M NaCl and had the same overall size (dimensions) as the bilayer-complex systems.

*Simulation procedure, software, and parameters.* All simulations were performed using GROMACS 2022.5 [27] with the Martini v.2 force field and the Elnedyn elastic network [28]. Following equilibration, three replica simulations were started for each bilayer-complex system by randomizing the initial atom velocities. The 7SC0 and 9DN0 replicas were run for 8 $\mu$s each, and the 9DN1 replicas for 14 $\mu$s each, all with a time step of 20 fs. A single replica simulation of 8 $\mu$s was run for the bilayer-only systems. A v-rescale thermostat and a semi-isotropic Parrinello-Rahman barostat were used with an update frequency of nsttcouple = nstpcouple = 10 steps. The Verlet cutoff scheme was used with a neighbor-list update frequency of nstlist = 25, an outer cutoff distance of rlist = 1.265 nm, and an inner cutoff of rl = 1.1 nm. All simulations were run at a constant temperature of 310.15 K. Furthermore, the Elnedyn elastic network used in the simulations with caveolin preserves the secondary structure of the complexes, but not their tertiary arrangement. To preserve the structure of the central β-barrel in particular, we applied a harmonic restraint between a pair of residues across all protomers: residues ASN173 and GLU177 in 7SC0, ILE143 and SER151 in 9DN0, and between LEU220 and GLU230 in 9DN1. This was done using the GROMACS pull code, with umbrella pulling at a pull rate of 0.0 nm/ps and a force constant of 1000 kJ $mol^{-1}$ $nm^{-1}$. These restraints have been shown not to have an effect on any measurable bilayer-caveolin interactions for 7SC0 [29].

*Simulations analysis.* All simulations were run until the protein structure was stabilized (as judged by the time evolution of the protein RMSD) for at least 4 $\mu$s. The final 1.5 $\mu$s of each trajectory was then used for analysis, with the frames output every 300 ps. For the bilayer-only systems, the analysis was performed on the last 2 $\mu$s of the trajectory.

*Mean curvature and lipid splay.* To calculate the mean curvature relative to the protein complex, we first centered and aligned the trajectory on the complex in the xy-plane. The positions of the PO4 beads of all lipids in the distal leaflet were then interpolated to obtain a surface on a grid with 3 Å grid spacing, and its mean curvature was calculated with *surfature* function in MATLAB R2025a. The lipid chain splay was calculated on the same aligned trajectory as the curvature. Two vectors were defined for each lipid between the C3 and the corresponding glycerol backbone bead of each chain. The splay angle between these two vectors was calculated for all lipids in the distal leaflet and mapped onto the same grid as the curvature for visualization. These calculations were done on each frame, and the results were averaged over all frames in the trajectory.

*Solvent accessible surface area (SASA).* The solvent-accessible surface area (SASA) was calculated using the GROMACS SASA tool. The radii of each bead in the system were specified based on the σ-parameter of the Lennard-Jones interactions of the beads, according to $r_{vdW} = \sqrt[6]{2} \cdot \sigma/2$ where $r_{vdW}$ is the van der Waals (vdW) radius of the bead, and σ is the σ-parameter of Lennard-Jones interactions between the bead of interest and other beads. The vdW radius of a water bead, 0.26 nm, was used as the probe size in the SASA tool.

*Protein complex hydrophobicity map.* To create a map of the distribution of hydrophobic residues in the 3D structure of each complex, we used the corresponding all-atom structures of the complexes. Consecutive slabs of 2 Å thickness were defined along an axis parallel to the beta barrel and passing through its center. Within each slab, we iterated over 2 Å-wide radial bands at increasing radii. The sum of the hydrophobicity values from the Kyte-Doolittle hydropathy scale [30] of each atom in each band was then averaged over all atoms in the band. The results were used to construct the hydrophobicity maps plotted in Figure 3H, where each grid point corresponds to a radial band.

*Tilt and thickness.* The tilt and thickness were calculated after centering each trajectory on the complex. For the tilt, a vector was defined from the geometric center of the C4 bead to the geometric centers of the PO4, GL1, and GL2 beads. The angle between this vector and the bilayer normal (i.e., the z-axis of the simulation box) was defined as the tilt of the lipid. The tilts were calculated iteratively, averaged over radial bands of width 3 Å starting from the protein boundary in each frame, and then averaged across all frames. The leaflet thickness was defined as the distance between the geometric centers of all PO4 and C4 beads within each of the 3 Å-wide radial bands and was calculated by averaging over all frames.

*Proximal leaflet lipid contacts.* Quantifying the interactions between protein residues and bilayer lipids was done by counting the number of contacts of a protein residue with any PO4, GL1, and GL2 bead. By calculating the radial density distribution function of the PO4, GL1, and GL2 beads relative to the protein, we defined a cutoff distance of 6 Å for a bead to be considered in contact with a protein residue. We then counted the number of contacts at the start and end of each simulation. The counts at the start were averaged over the first 5 ns, and the counts at the end were averaged over the final 1.5 µs.

*Phosphate min/max position difference.* The difference between the maximum and minimum position of all phosphates was calculated directly from the bilayer-only systems, or after centering each complex-bilayer trajectory on the complex. For all trajectories, the difference between the maximum and minimum z-positions of any PO4 beads in both leaflets was calculated in every frame. The differences were then plotted as a histogram. For the complex bilayer systems, the differences were also averaged over the three replicas.

*Mean curvature difference.* To compare the mean curvature between the complex-bilayer systems and the bilayer-only systems, a moving average of the mean curvature was calculated. First, the mean curvature was calculated for each system as detailed above. A window of dimension $(2j+1) \times (2j+1)$, where $j$ is the number of grid points, was convolved with the mean curvature matrix in every frame, after which the average mean curvature of each window was calculated. These averages were then summed to obtain the total mean curvature in the current frame; the total mean curvatures over the trajectory were then plotted as a histogram. To determine the optimal value of $j$, we systematically varied the parameter in the 7SC0 complex-bilayer system. The value that gave the largest total mean curvature was $j = 7$ which was then used for the analysis of all systems (hence the window dimension was $15 \times 15$ grid points).

### Supplementary Note 1

### Continuum mechanics model

We present a minimal model designed to investigate how caveolins interact with lipids and contribute to curvature generation during caveolae formation. Our model concerns a disc-shaped caveolin complex with radius $R_0$ embedded in a lipid monolayer by displacing lipids from the corresponding leaflet. We denote the local thickness of the monolayer by the scale field $u(\boldsymbol{r})$, where $\boldsymbol{r} = (x, y)$. The equilibrium thickness of the unperturbed monolayer is denoted by $l_0$. The elastic energy associated with the thickness deformation of the lipid leaflet can then be written in the form [31-34]

$$F = \int \left[\frac{\kappa}{2}(\nabla^2 u)^2 + \frac{\sigma}{2}(\nabla u)^2 + \frac{K_{\mathrm{A}}}{2}\left(\frac{u - l_0}{l_0}\right)^2\right] \mathrm{d}A$$

where $\kappa$ is the bending modulus, $\sigma$ is the membrane tension, and $K_{\mathrm{A}}$ is the compression modulus or stiffness of the monolayer associated with thickness deformation. The first two terms in the integrand represent the energetic cost of bending and stretching the monolayer. The last term accounts for the energy penalty due to local compression or expansion of the lipid leaflet thickness relative to its equilibrium value.

Following the calculus of variations [31-34], one can obtain the Euler-Lagrange equation in the following form

$$\kappa\nabla^4 u - \sigma\nabla^2 u + \frac{K_{\mathrm{A}}}{l_0^2}u - \frac{K_{\mathrm{A}}}{l_0} = 0$$

The associated boundary conditions are specified as follows: $u(r = R_0) = L$, $u(r \to \infty) = l_0$, $u'(r = R_0) = s = \tan\theta$, $u'(r \to \infty) = 0$, where $s$ is the contact slope at the caveolin-monolayer contact interface and $\theta$ is the angle between the tangent to the monolayer and the horizontal at the contact point.

Solutions to this Euler-Lagrange equation can be found in [34-36] and can be summarized as

$u(r) = C_1\mathrm{K}_0(\lambda_1 r) + C_2\mathrm{K}_0(\lambda_2 r) + l_0,$

where $C_1$ and $C_2$ are the coefficients which are determined by the boundary conditions, $\mathrm{K}_0$ is the modified zero-order Bessel function of the second kind. Here, $\lambda_{1,2}$ are given by $\lambda_{1,2} = \sqrt{\frac{(\sigma/\kappa) \pm \sqrt{(\sigma/\kappa)^2 - 4K_{\mathrm{A}}/(\kappa l_0^2)}}{2}}$. By using the boundary conditions, the coefficients $C_1$ and $C_2$ are obtained as

$$C_1 = \frac{\lambda_2(L - l_0)\mathrm{K}_1(\lambda_2 R_0) + s\mathrm{K}_0(\lambda_2 R_0)}{\lambda_2\mathrm{K}_0(\lambda_1 R_0)\mathrm{K}_1(\lambda_2 R_0) - \lambda_1\mathrm{K}_0(\lambda_2 R_0)\mathrm{K}_1(\lambda_1 R_0)}$$

$$C_2 = \frac{\lambda_1(L - l_0)\mathrm{K}_1(\lambda_1 R_0) + s\mathrm{K}_0(\lambda_1 R_0)}{\lambda_1\mathrm{K}_0(\lambda_2 R_0)\mathrm{K}_1(\lambda_1 R_0) - \lambda_2\mathrm{K}_0(\lambda_1 R_0)\mathrm{K}_1(\lambda_2 R_0)}$$

For the data shown in Figure 3D, results were obtained by finding the thickness profile that minimizes the total energy. At a fixed value of $L$, we numerically solve the thickness profile for various contact slope $s$ and subsequently compute the associated energy. By identifying the value of $s$ that minimizes the total energy, we determine the optimal slope at the caveolin–monolayer interface and the thickness profile of the monolayer that corresponds to the energy minimum.

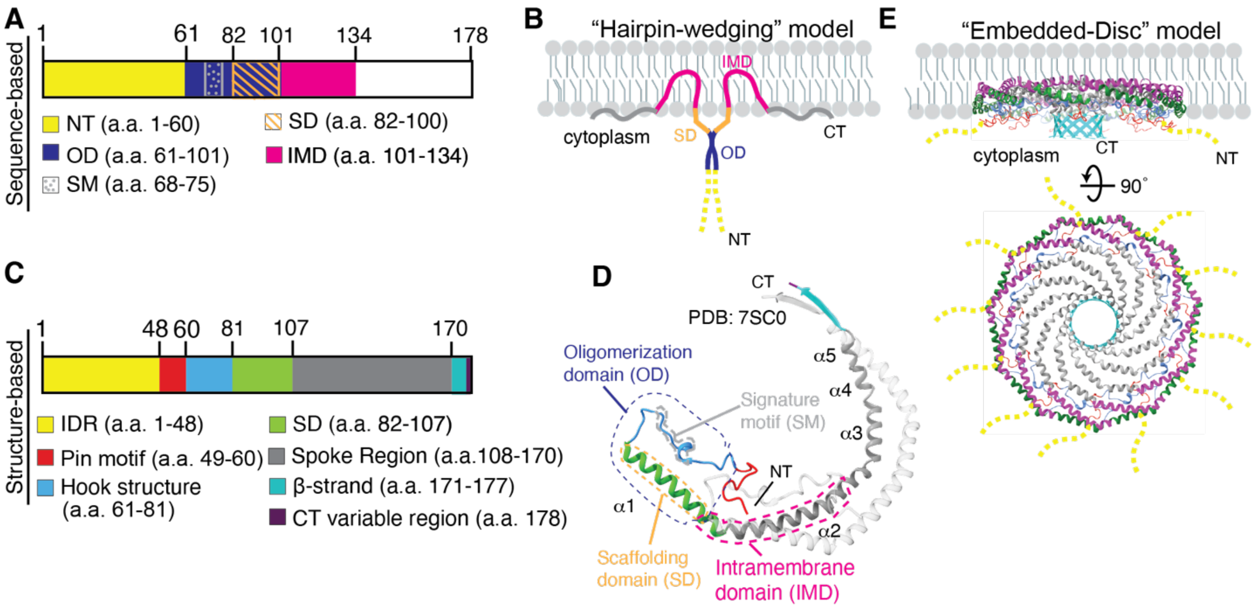

**Figure S1. Cryo-EM structure of *Hs* CAV1 presents a new model of domain architecture and function.** (**A**) Cartoon diagram of *Hs* CAV1 showing domain organization predicted using sequence-based analysis. NT: N-terminal region; OD: oligomerization domain; SM: signature motif; SD: scaffolding domain, IMD: intramembrane domain. (**B**) The "hairpin-wedging" model predicts that the IMD forms a "hairpin-like" structure that wedges into the membrane. NT: N-terminus; CT: C-terminus. Colors are same as panel A, except the undefined region of the complex (a.a. 134-178) is grey. (**C**) Cartoon diagram of *Hs* CAV1 showing domain organization from structure-based analysis. IDR: intrinsic disordered region. (**D**) Secondary structure representation of two *Hs* CAV1 protomers, with one colored the same as panel C. "Domains" annotated before the structure was available are circled in dashed lines. The NT IDR was not visible in the cryo-EM map. (**E**) The structure-based "embedded disc" model predicts that the entire disc sits within the bilayer. *Hs* CAV1 complex (PDB: 7SC0) is rotated 90˚ around the x-axis to show the surface of the embedded disc. Residues in the IDR (residues1-48) were not visible in the density map and are shown as dashed lines. Complex is colored as shown in panel C, with the exception that the IMD residues (residues 101-134) that span regions of the SD and SR are colored magenta.

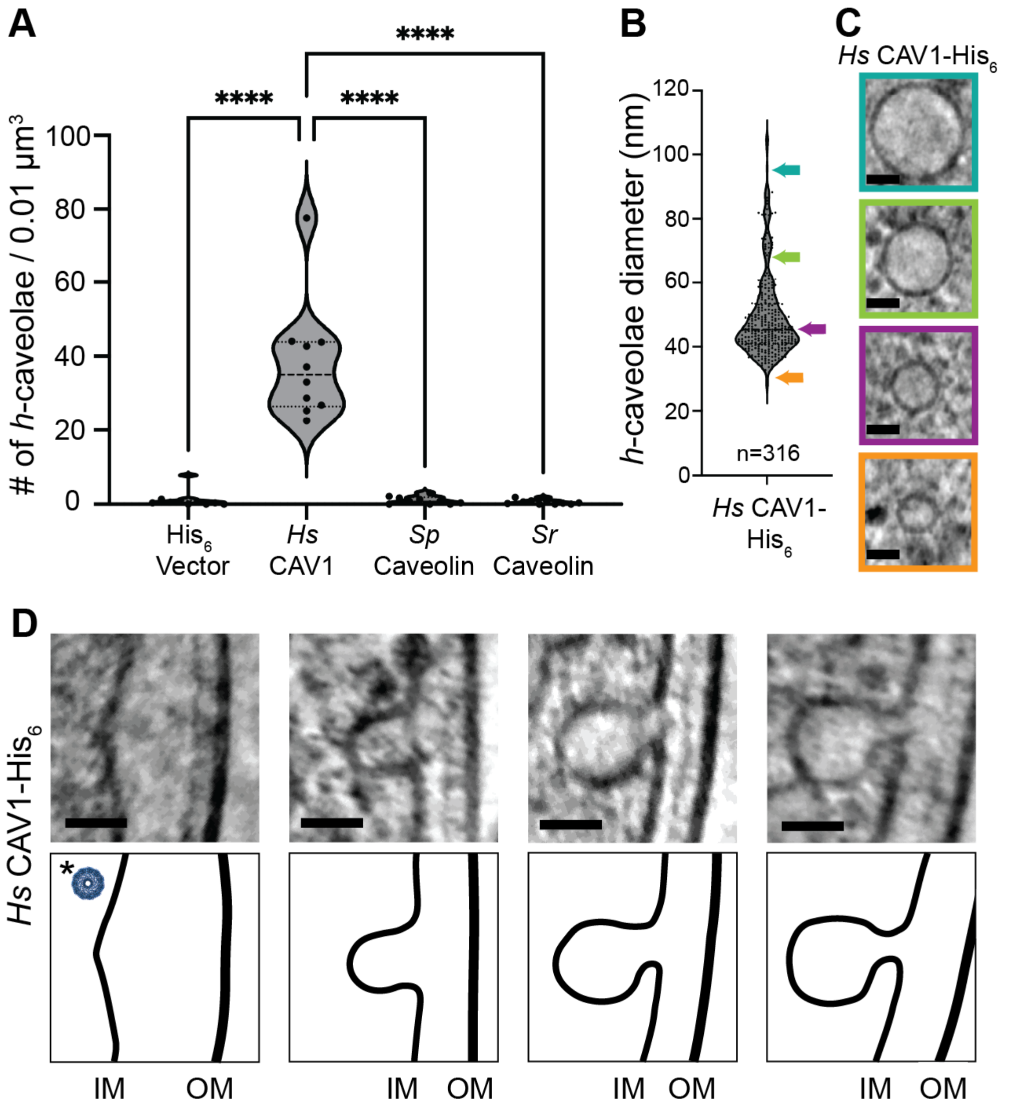

**Figure S2. Cryo-ET analysis of *E. coli* expressing Human (*Hs*), *S. purpuratus* (*Sp*), or *S. rosetta* (*Sr*) Caveolins.** (**A**) A violin plot showing the number of *h*-caveolae found per 0.01 µm$^3$ of cellular volume in *E. coli* expressing the His$_6$ vector, *Hs* CAV1-His$_6$, *Sp* Caveolin-His$_6$, or *Sr* Caveolin-His$_6$. **** *P* = <0.0001. (**B**) A violin plot of the diameter distribution of *h*-caveolae measured in 8 tomograms of *E. coli* expressing *Hs* CAV1-His$_6$. N=316 *h*-caveolae measured for diameter. Colored arrows indicate the diameter of the *h*-caveolae shown in panel C. (**C**) Representative images of various-sized *h*-caveolae found in tomograms of *E. coli* expressing *Hs* CAV1-His$_6$. Images are the sum of 13 slices. Color of outline corresponds to arrows shown in panel B. Scale bars, 30 nm. (**D**) Representative images of the bacterial inner membrane in tomograms of vitrified *E. coli* expressing *Hs* CAV1-His$_6$ (top) and the corresponding cartoon outline (bottom). Images in the top panel are the sum of 13 slices. *, Cryo-EM density map of the *Hs* CAV1 complex (blue, EMDB: 25007) is shown to scale with the traced bacterial membranes in the bottom panel. IM, inner membrane; OM, outer membrane. Scale bars, 30nm.

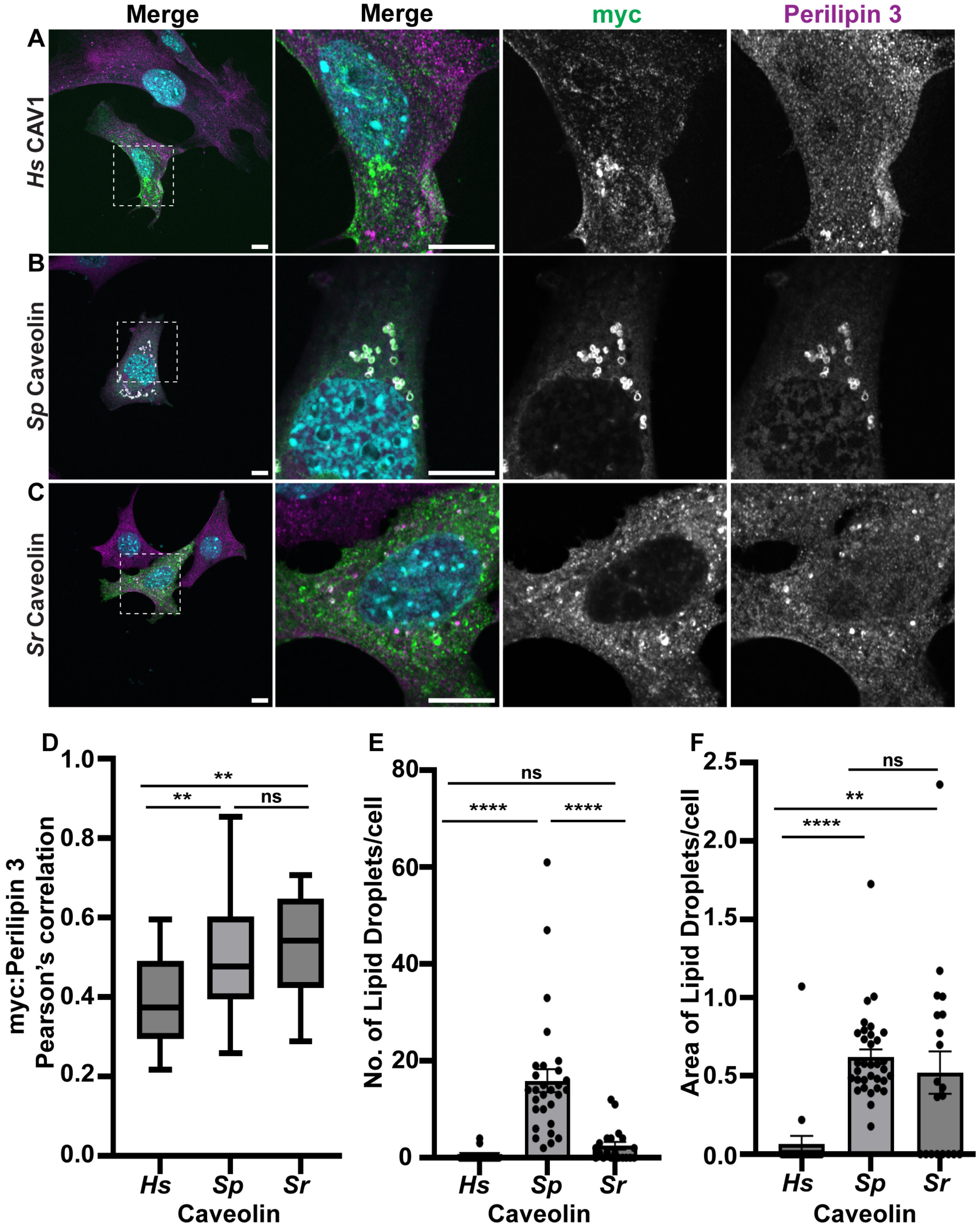

**Figure S3. *Sp* and *Sr* Caveolins expressed in mammalian cells are primarily localized to lipid droplets.** (**A-C**) Representative confocal images shown for CAV1$^{-/-}$ MEF cells expressing myc-tagged *Hs* CAV1, *Sp* Caveolin, or *Sr* Caveolin. Transfected cells expressing the indicated constructs for 24 h were fixed and immunostained with antibodies against myc (green) and the lipid droplet marker Perilipin 3 (magenta). Dashed boxes in the left column indicate the position of zoomed images shown in the other three columns. Scale bar, 10 µm. (**D**) Pearson's Correlation analysis (*Hs* CAV1, n=30, *Sp* Caveolin, n=20, *Sr* Caveolin, n=20 ROIs from at least two independent experiments were used to determine the colocalization of caveolins and Perilipin 3. (**E**) Number of lipid droplets per cell under each condition was counted manually using the counter plugin in Fiji. (**F**) The number of lipid droplets per cell under each condition was calculated by outlining individual lipid droplets, and the area of each droplet was determined using the measurement function in Fiji. One-way ANOVA with Tukey's test (≥3 groups) was used to calculate p-values. **, $p < 0.001$; ****, $p < 0.0001$; n.s., not significant.

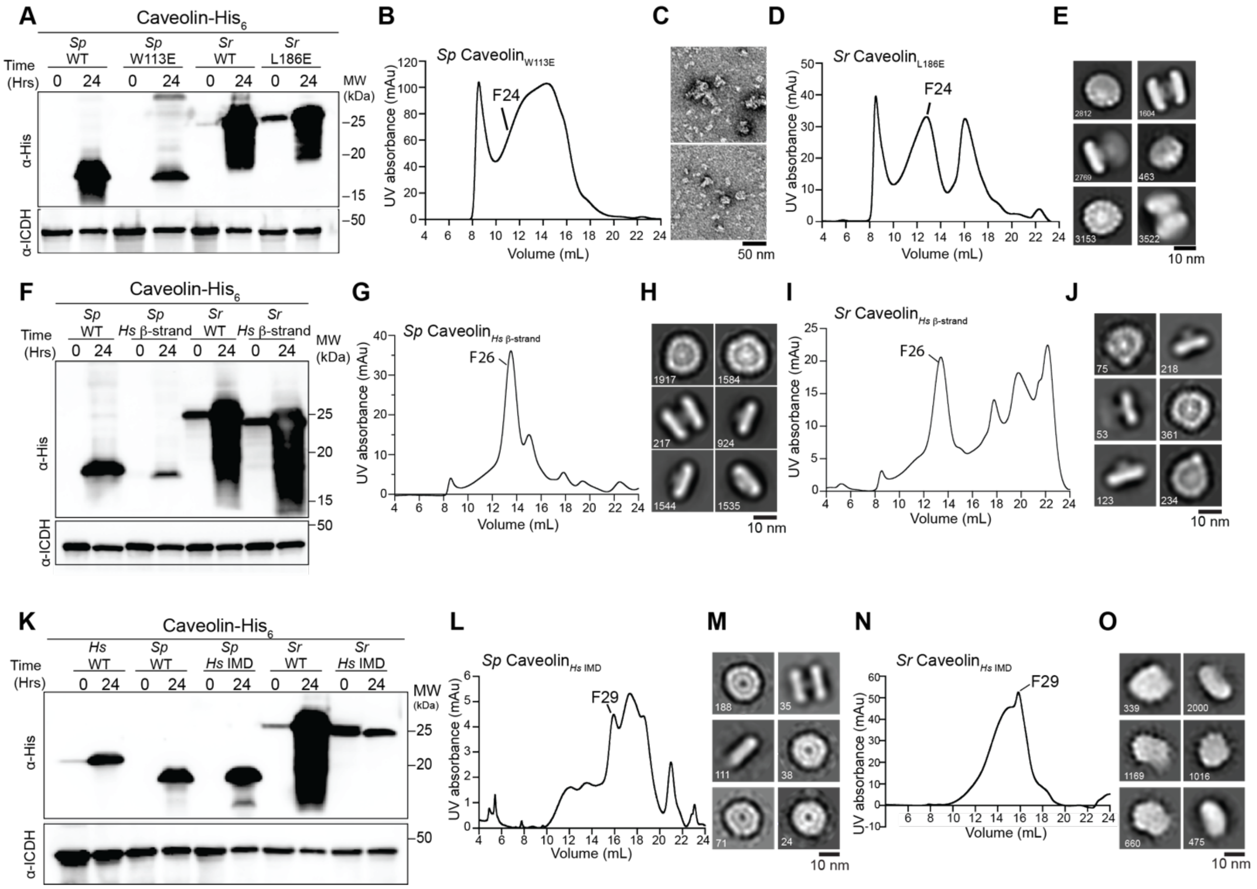

**Figure S4. Biochemical characterization of *Sp* and *Sr* Caveolin mutants.** (**A**) Western blot analysis showing expression of His$_6$-tagged *Sp* Caveolin$_{W113E}$ or *Sr* Caveolin$_{L186E}$ in *E. coli*. (**B**) FPLC trace of *Sp* Caveolin$_{W113E}$ purification. Line marks fraction used for negative stain analysis shown in panel C. (**C**) Representative micrograph of negatively stained *Sp* Caveolin$_{W113E}$ purification. Scale bar, 50 nm. (**D**) FPLC trace of *Sr* Caveolin$_{L186E}$. The fraction used for negative stain analysis, shown in panel E, is marked by a line. (**E**) 2D classes of negatively stained *Sr* Caveolin$_{L186E}$ complexes. Scale bar, 10 nm. The number of particles in each class is indicated. (**F**) Western blot analysis showing expression of His$_6$-tagged *Sp* Caveolin$_{Hs\ \beta\text{-strand}}$ or *Sr* Caveolin$_{Hs\ \beta\text{-strand}}$ in *E. coli*. (**G**) FPLC trace of *Sp* Caveolin$_{Hs\ \beta\text{-strand}}$ purification. Line marks fraction used for negative stain analysis shown in panel H. (**H**) 2D classes of negatively stained *Sp* Caveolin$_{Hs\ \beta\text{-strand}}$ complexes. Scale bar, 10 nm. Number of particles in each class indicated. (**I**) FPLC trace of *Sr* Caveolin$_{Hs\ \beta\text{-strand}}$ purification. Arrow marks fraction used for negative stain analysis shown in panel J. (**J**) 2D classes of negatively stained *Sr* Caveolin$_{Hs\ \beta\text{-strand}}$ complexes. Scale bar, 10 nm. The number of particles in each class is indicated. (**K**) Western blot analysis showing expression of His$_6$-tagged *Sp* Caveolin$_{Hs\ IMD}$ or *Sr* Caveolin$_{Hs\ IMD}$ in *E. coli*. (**L**) FPLC trace of *Sp* Caveolin$_{Hs\ IMD}$ purification. Line marks fraction used for negative stain analysis shown in panel M. (**M**) 2D classes of negatively stained *Sp* Caveolin$_{Hs\ IMD}$ complexes. Scale bar, 10 nm. The number of particles in each class is indicated. (**N**) FPLC trace of *Sr* Caveolin$_{Hs\ IMD}$ purification. The fraction used for negative-stain analysis, shown in panel O, is marked with a line. (**O**) 2D classes of negatively stained *Sr* Caveolin$_{Hs\ IMD}$ complexes. Scale bar, 10 nm. The number of particles in each class is indicated. (A, F, and K) Membranes were probed with anti-His (α-His) or anti-ICDH (α-ICDH) antibodies. (E, H, J, O) The number of particles found in each average is shown in the corner.

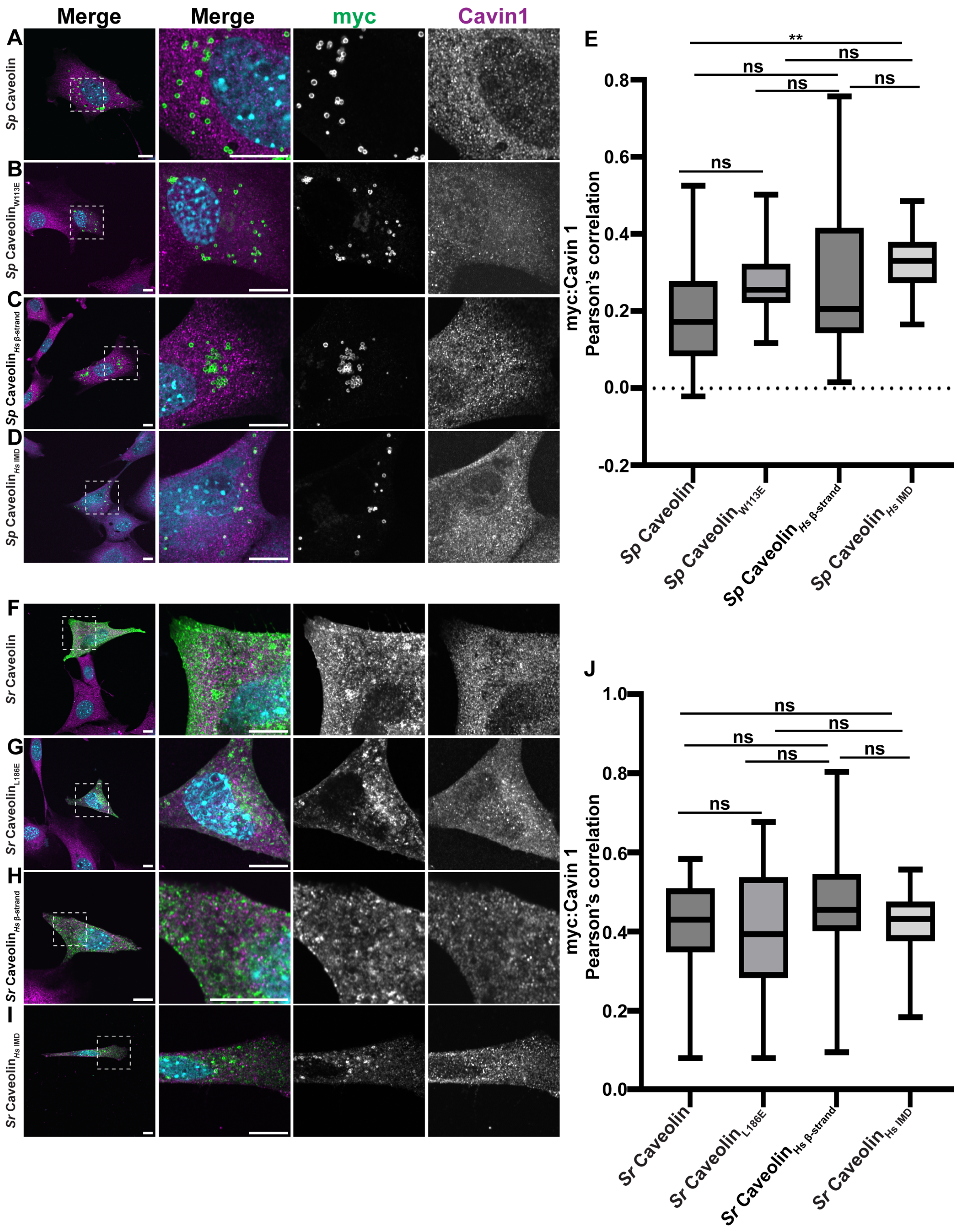

**Figure S5. Analysis of *Sp* and *Sr* Caveolin mutants in CAV1$^{-/-}$ MEFs. (A-D)** Representative confocal images shown for CAV1$^{-/-}$ MEF cells expressing myc-tagged *Sp* Caveolins: *Sp* Caveolin (A), *Sp* Caveolin$_{W113E}$ (B), *Sp* Caveolin$_{Hs\ \beta\text{-strand}}$ (C), or *Sp* Caveolin$_{Hs\ IMD}$ (D). (**E**) Graph showing Pearson's correlation analysis for determining colocalization of *Sp* Caveolin and Cavin1 in panels A-D. (**F-I**) Representative confocal images shown for CAV1$^{-/-}$ MEF cells expressing myc-tagged

*Sp* Caveolins: *Sr* Caveolin (F), *Sr* Caveolin$_{L186E}$ (G), *or Sr Caveolin$_{Hs\ \beta\text{-strand}}$* (H), or *Sr* Caveolin$_{Hs\ IMD}$ (I). (**J**) Graph showing Pearson's correlation analysis for determining colocalization of *Sr* Caveolin and Cavin1 in panels F-I. (A-D and F-I) Transfected cells expressing indicated constructs for 24 h, were fixed, and immunostained with anti-myc (green) and Cavin1 (magenta) antibodies. Dashed boxes in left column (scale bars, 10 µm) indicate the position of zoomed images shown in the other columns. Scale bars, 10 µm. For Pearson's correlation analysis *Sp* Caveolin, n=30; *Sp* Caveolin$_{W113E}$, n=21; *Sp* Caveolin$_{Hs\ \beta\text{-strand}}$, n=30; *Sp* Caveolin$_{Hs\ IMD,}$ n=20; *Sr* Caveolin$_{,}$ n=25; *Sr* Caveolin$_{L186E,}$ n=20; *Sr* Caveolin$_{Hs\ \beta\text{-strand}}$, n= 24; *Sr* Caveolin$_{Hs\ IMD,}$ n=18. ROIs were used from ≥ 2 independent experiments. A one-way ANOVA with Tukey's test was used to calculate p-values. **, $p < 0.01$; n.s., not significant.

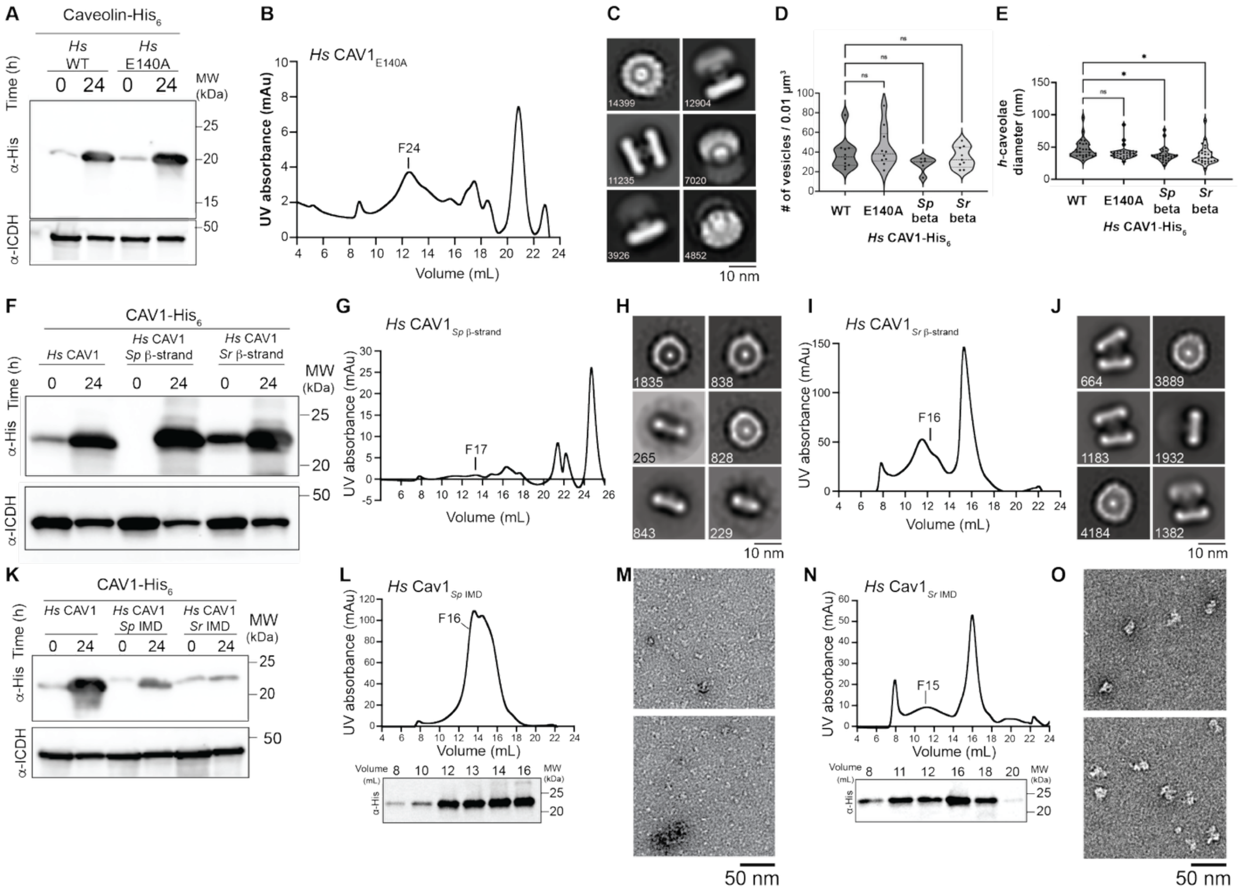

**Figure S6. Biochemical characterization of *Hs* CAV1 mutants.** (**A**) Western blot analysis showing expression of $His_6$-tagged *Hs* CAV1 and $CAV1_{E140A}$ in *E. coli*. (**B**) FPLC trace of *Hs* $CAV1_{E140A}$. Line marks fraction used for negative stain analysis shown in panel C. (**C**) 2D class averages of purified and negative stained *Hs* $CAV1_{E140A}$ complexes. Scale bar, 10 nm. The number of particles in each class is indicated. (**D**) A violin plot showing the number of *h*-caveolae found per 0.01 μm$^3$ of cellular volume for *E. coli* expressing $His_6$-tagged *Hs* CAV1, $CAV1_{E140A}$, $CAV1_{Sp\ \beta\text{-strand}}$, or $CAV1_{Sr\ \beta\text{-strand}}$. Number of tomograms analyzed include: 10 for *Hs* CAV1, 10 for *Hs* $CAV1_{E140A}$, 6 for *Hs* $CAV1_{Sp\ \beta\text{-strand}}$, and 9 for *Hs* $CAV1_{Sr\ \beta\text{-strand}}$. ANOVA tests, ns $P$ = 0.7149, 0.4361, 0.9186. (**E**) Violin plot showing *h*-caveolae diameters measured in tomograms of *E. coli* expressing $His_6$-tagged *Hs* CAV1, $CAV1_{E140A}$, $CAV1_{Sp\ \beta\text{-strand}}$, or $CAV1_{Sr\ \beta\text{-strand}}$. 6 tomograms per sample were analyzed. ANOVA tests, ns $P$ = 0.2416, * $P$ = 0.0258, 0.0122. (**F**) Western blot analysis showing expression of $His_6$-tagged *Hs* CAV1, $CAV1_{Sp\ \beta\text{-strand}}$, or $CAV1_{Sr\ \beta\text{-strand}}$ in *E. coli*. (**G**) FPLC trace of *Hs* $CAV1_{Sp\ \beta\text{-strand}}$. Line marks fraction used for negative stain analysis shown in panel H. (**H**) 2D class averages of purified and negative stained $CAV1_{Sp\ \beta\text{-strand}}$ complexes. Scale bar, 10 nm. The number of particles in each class is indicated. (**I**) FPLC trace of *Hs* $CAV1_{Sr\ \beta\text{-strand}}$. Line marks fraction used for negative stain analysis shown in panel J. (**J**) 2D class averages of purified and negative stained $CAV1_{Sr\ \beta\text{-strand}}$ complexes. Scale bar, 10 nm. The number of particles in each class is indicated. (**K**) Western blot analysis showing expression of $His_6$-tagged *Hs* CAV1, $CAV1_{Sp\ IMD}$, or $CAV1_{Sr\ IMD}$ in *E. coli*. (**L**) FPLC trace of *Hs* $CAV1_{Sp\ IMD}$. Anti-His Western blot of FPLC fractions is shown below the trace. Line marks fraction used for negative stain analysis shown in panel M. (**M**) Two representative images of the of purified and negative stained $CAV1_{Sp\ IMD}$ purification. Scale bar, 50 nm. (**N**) FPLC trace of *Hs* $CAV1_{Sr\ IMD}$. Anti-His Western blot of each FPLC fraction shown below the trace. Arrow marks fraction used for negative stain analysis shown in panel O. (**O**) Two representative images of the purified and negative stained CAV1$_{Sr\ \mathrm{IMD}}$ purification. Scale bar, 50 nm. (A, F, K, L, and N) For Western blots the membranes were probed with anti-His ($\alpha$-His) or anti-ICDH ($\alpha$-ICDH) antibodies. (C, H, and J) The number of particles found in each average is shown in the corner.

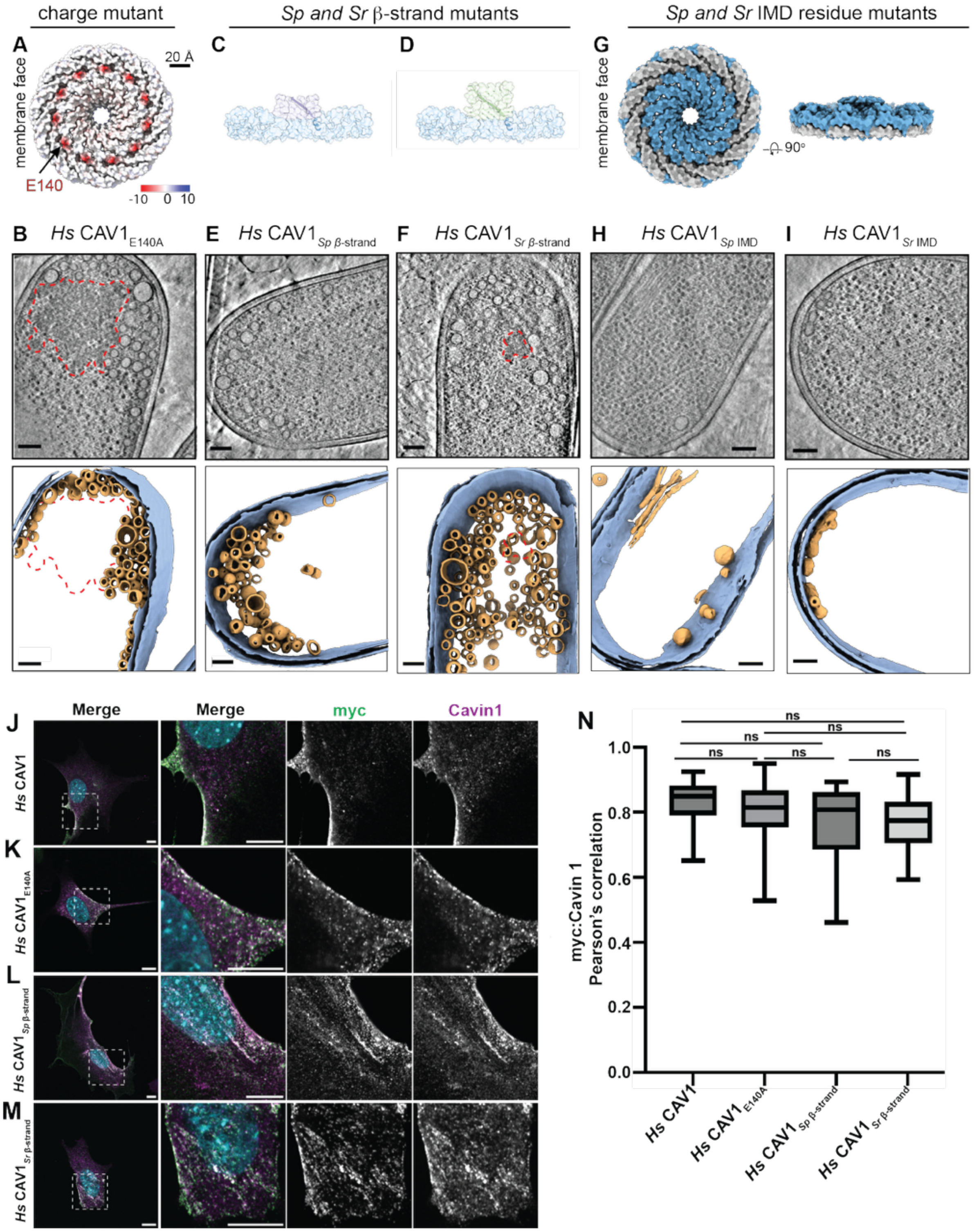

**Figure S7. Analysis of *h*-caveolae and caveolae formation of *Hs* CAV1 mutants.** (**A**) Space-filling model showing the charge distribution on the hydrophobic face of the *Hs* CAV1 complex. The arrow points to Glu140. (**B**) Slices through representative deconvoluted tomogram (top panel) and corresponding segmentation (bottom panel) of *E. coli* expressing $His_6$-tagged *Hs* $CAV1_{E140A}$. (**C, D**) Cut through of space-filling models of the *Hs* $CAV1_{Sp\ \beta\text{-strand}}$ or *Hs* $CAV1_{Sr\ \beta\text{-strand}}$ mutants with one β-strand shown as a ribbon diagram. (**E, F)** Slices through representative deconvoluted tomograms (top panel) and corresponding segmentations (bottom panel) of *E. coli* expressing $His_6$-tagged *Hs* Cav1$_{Sp\ \beta\text{-strand}}$ (E) or *Hs* CAV1$_{Sr\ \beta\text{-strand}}$ (F). (**G**) Surface model of the predicted membrane facing surface of *Hs* CAV1 rotated 90˚ around the x-axis. The position of the *Sp* or *Sr* IMD residues in the mutants is shown in grey. (**H-I**) Slices through representative deconvoluted tomograms (top panel) and corresponding segmentations (bottom panel) of *E. coli* expressing $His_6$-tagged *Hs* CAV1 containing IMD residues from *Sp* or *Sr* Caveolins. (B, E, F, H, and I) Bacterial membranes are shown in blue, *h*-caveolae or other small vesicles are shown in gold. Red dashed lines outline electron-dense regions that could represent inclusion bodies. Scale bars, 100 nm. (**J-M**) Representative confocal microscopy images shown for CAV1$^{-/-}$ MEF cells expressing myc-tagged *Hs* CAV1 (J), *Hs* CAV1$_{E140A}$ (K), *Hs* CAV1$_{Sp\ \beta\text{-strand}}$ (L), and *Hs* CAV1$_{Sr\ \beta\text{-strand}}$ (M). Transfected cells expressing the indicated constructs for 24 h were fixed and immunostained with anti-myc (green) and Cavin1 (magenta) antibodies. Dashed boxes in the left column (scale bar, 10 μm) indicate the position of zoomed images shown in the other columns. Scale bar, 10 μm. (**N**) Colocalization of caveolin and Cavin1 was determined by Pearson's Correlation analysis (*Hs* CAV1, n=20; *Hs* CAV1$_{E140A}$, n=35; *Hs* CAV1$_{Sp\ \beta\text{-strand}}$, n=38; *Hs* CAV1$_{Sr\ \beta\text{-strand}}$, n=4. ROIs from ≥ 2 independent experiments). One-way ANOVA with Tukey's test (≥3 groups) was used to calculate p-values. n.s., not significant.

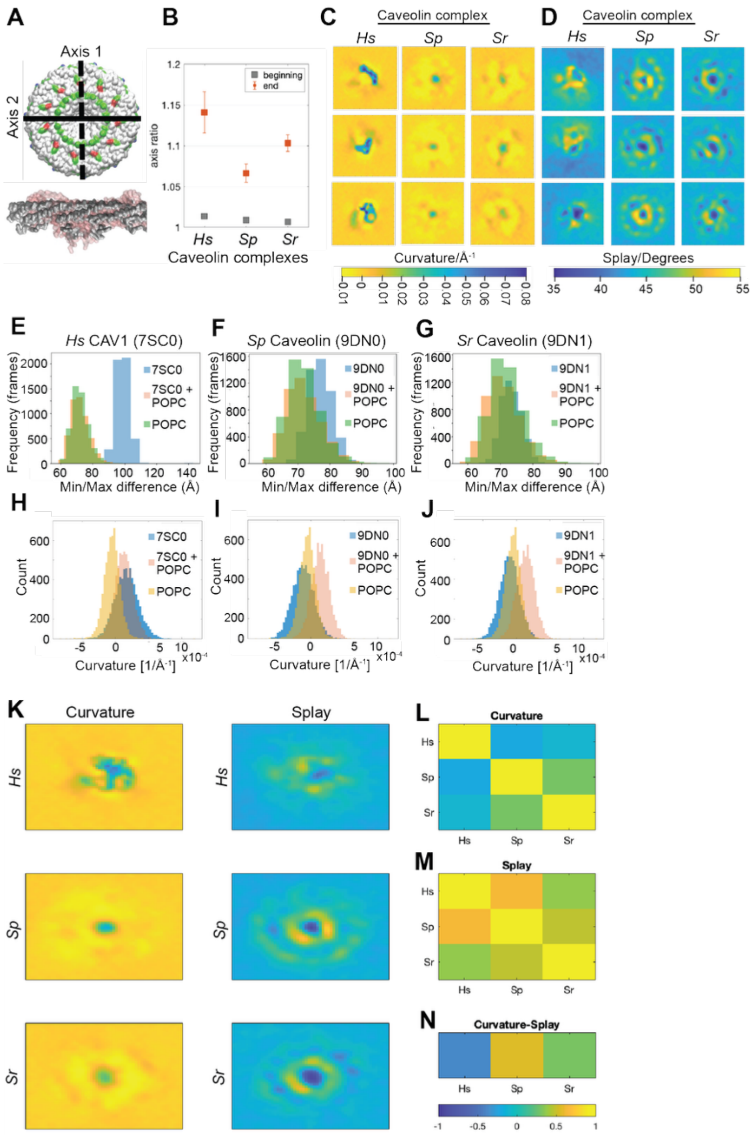

**Figure S8. Analysis of complex shape, membrane shape, and lipid conformation in simulations.** (**A-B**) Flexibility of the caveolin complexes observed in simulations. (A) The Axis 1 and 2 are labeled using the structure of the *Hs* CAV1 complex (7SC0), with residues colored by hydrophobicity: polar green, basic blue, acidic red, and nonpolar white (top panel). Starting structure (grey) and ending structure (salmon) overlayed (bottom panel). (B) Aspect ratio of the two axes of the structure for each complex calculated at the beginning and end of the simulation for each caveolin complex. Errors represent the standard deviation across the three replica simulations. (**C-D**) Mean curvature (C) and lipid splay angles (D) calculated in the distal leaflet of three independent replica simulations (top to bottom) of *Hs* CAV1 (left), *Sp* Caveolin (middle), and *Sr* Caveolin (right) embedded in a POPC membrane. The membrane in the *Hs* CAV1 trajectories has an overall higher mean curvature relative to the bilayer in the *Sp* and *Sr* Caveolin complex simulations (C), while no consistent trends are observed in lipid splay between the three complexes (D). The small central region showing higher curvature and lower splay for *Sp* and *Sr* Caveolin complexes is due to a few lipids being pulled towards the hydrophobic interior region of the central β-barrel, ordering the lipid tails in a more parallel manner. All plots are 165 × 165 Å and centered on the caveolin complex. (**E**–**G**) Largest distance (along z-axis) between lipid headgroups (PO4 beads in MARTINI notation) in every trajectory frame for *Hs* CAV1 (E), *Sp* Caveolin (F) and *Sr* Caveolin (G) complexes plotted as a histogram. The complex bilayer system is in blue. In addition, we simulated a lipid-only bilayer (without protein) with either equal numbers of lipids in both leaflets (balanced POPC) or with the same number of lipids removed from one leaflet as in the corresponding complex-bilayer simulation (imbalanced POPC) in green and orange, respectively. (**H-J**) Window averaging of the mean curvature for *Hs* CAV1 (H), *Sp* Caveolin (I), and *Sr* Caveolin (J) complexes from individual trajectory frames plotted as a histogram (see methods). The complex bilayer system is shown in blue, and the corresponding imbalanced and balanced POPC systems are shown in pink and yellow, respectively. (**K**) Average curvature (left) and splay (middle) maps for *Hs* CAV1 complex (top), *Sp* Caveolin complex (middle), and *Sr* Caveolin complex (bottom) calculated from the three individual replica simulations for each system. The color scales are the same as those in panels C and D. (**L-N**) Pairwise correlation between the curvature (L) and splay (M) maps is shown, as well as the correlation between curvature and splay for each system (N). All correlations are Spearman correlations with a color scale as indicated on the plot.

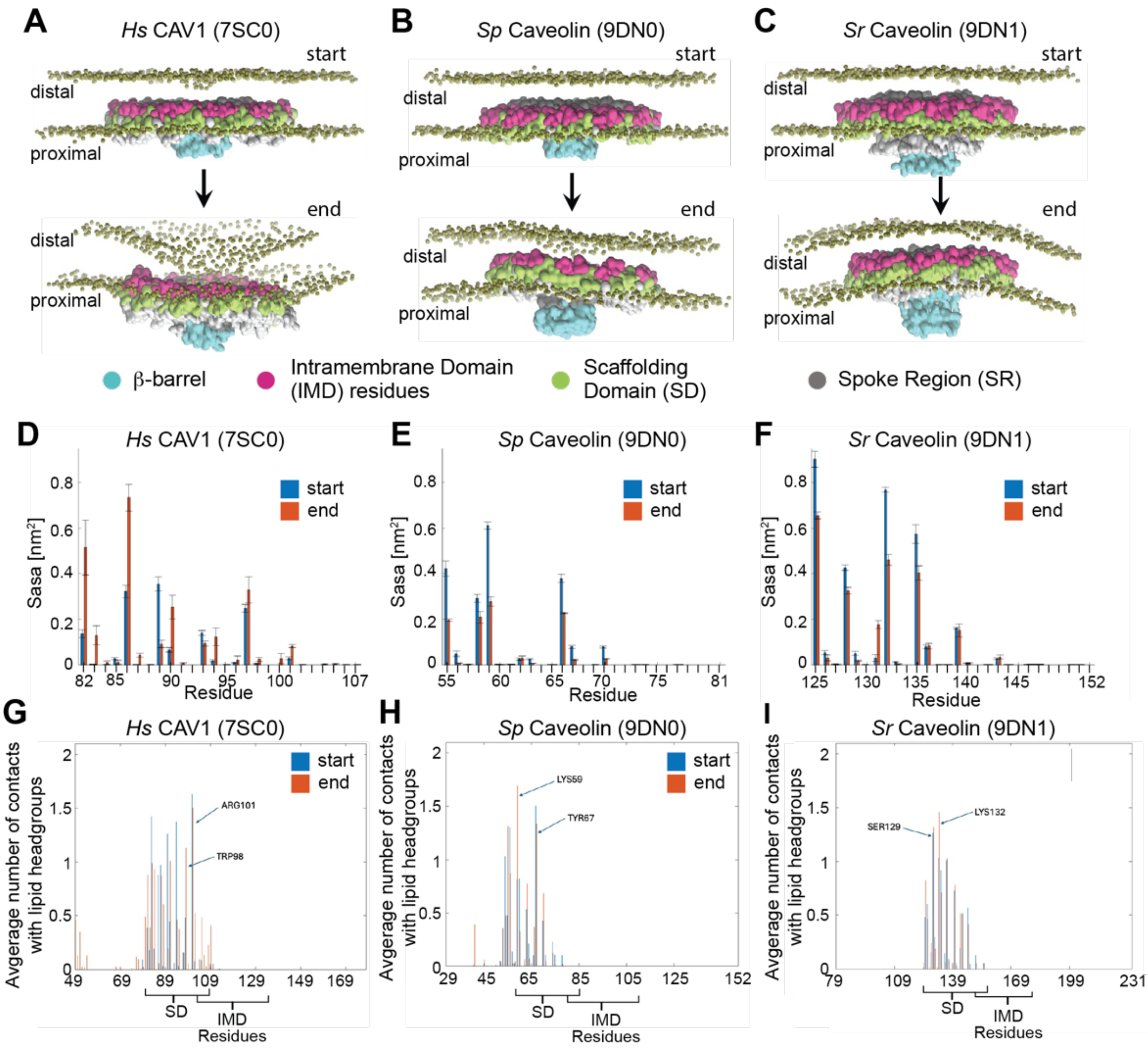

**Figure S9. Analysis of protein-lipid interactions in simulation trajectories.** (**A-C**) Simulations snapshots from the start and end of the simulations for *Hs* CAV1 (A), *Sp* Caveolin (B), and *Sr* Caveolin (C) complexes. The complexes are shown in a quicksurf surface representation. β-barrels shown in green, scaffolding domain (SD) shown in blue, intramembrane domain (IMD) residues shown in red, and shared residues between the SD and IMD residues shown in yellow. (**D-F**) Solvent accessible surface area (SASA) at the start (blue) and end (red) of the simulations for the residues belonging to the scaffolding domain (SD) regions of *Hs* CAV1 (D), *Sp* Caveolin (E), and *Sr* Caveolin (F) complexes. The increase in CSD residue exposure for the *Hs* CAV1 complex, but not for *Sp* and *Sr* Caveolin complexes, is consistent with proximal lipids sliding along the outer rim of the complex toward the bilayer interior. (**G-I**) Proximal leaflet lipid contacts at the start (blue) and end (red) of the simulations for the *Hs* CAV1 (G), *Sp* Caveolin (H) and *Sr* Caveolin (I) complexes. The two residues for each complex with highest number of lipid contacts at the end of the simulations are labeled. The residues belonging to the IMD region of *Hs* CAV1 have increased lipid contacts toward the end of the simulation, which is not seen for *Sp* and *Sr* Caveolins.

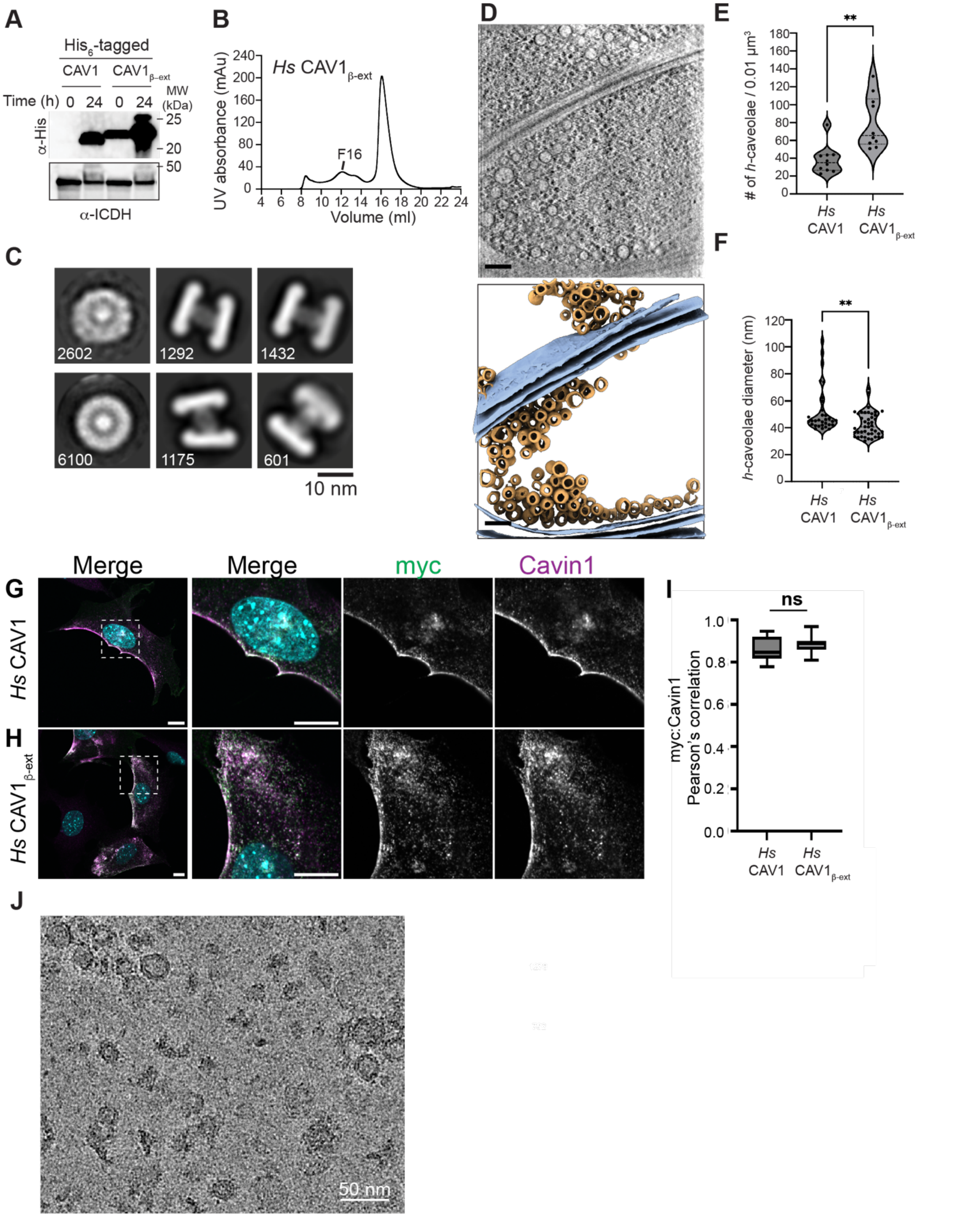

**Figure S10. Characterization of *Hs* CAV1 $_{\beta\text{-ext}}$.** (**A**) Western blot analysis showing expression of His$_6$-tagged *Hs* CAV1 or *Hs* CAV1$_{\beta\text{-ext}}$ (*Hs* CAV1 with an extended β-strand encoding *Sr* residues 225-233) in *E. coli*. Membranes probed with anti-His (α-His) or anti-ICDH (α-ICDH) antibodies. (**B**) FPLC trace of *Hs* CAV1 $_{\beta\text{-ext}}$ purification. Arrow marks fraction used for negative stain analysis shown in panel C. (**C**) Representative 2D classes of negatively stained *Hs* CAV1 $_{\beta\text{-ext}}$ complexes. Scale bar, 10 nm. (**D**) Cryo-ET analysis of *E. coli* expressing $His_6$-tagged *Hs* CAV1 $_{\beta\text{-ext}}$. Slice through a representative deconvoluted tomogram (top panel) and its corresponding segmentation (bottom panel). Bacterial membranes are shown in blue, and *h*-caveolae are shown in gold. Scale bars, 100 nm. (**E, F**) Violin plots showing analysis of the number of *h*-caveolae present per 0.01 $\mu m^3$ cell volume (E) or diameters of vesicles visualized in tomograms (F) of *E. coli* expressing $His_6$-tagged *Hs* CAV1 or *Hs* CAV1 $_{\beta\text{-ext}}$. For data in panel E, 10 tomograms were analyzed for *Hs* CAV1$_{\beta\text{-ext}}$. Paired t-test, $^{**}P = 0.0036$. For data in panel F, 10 tomograms were analyzed for *Hs* CAV1 $_{\beta\text{-ext}}$. Unpaired t-test, $^{**}P = 0.0022$. Data for *Hs* CAV1 in panels E and F are the same as shown in Figure S6D and E (analysis of 10 tomograms). (**G, H**) Representative confocal images shown for CAV1$^{-/-}$ MEF cells expressing myc-tagged *Hs* CAV1 (G) or *Hs* CAV1 $_{\beta\text{-ext}}$. (H). Transfected cells expressing the indicated constructs for 24 h were fixed and immunostained with anti-myc (green) and Cavin1 (magenta) antibodies. Dashed boxes in the left column (scale bars, 10 µm) indicate the position of zoomed images shown in the other columns. Scale bars, 10 µm. (**I**) Graph showing Pearson's correlation analysis for determining colocalization of caveolin and Cavin1 in G and H. For Pearson's correlation analysis, *Hs* CAV1, n=10, *Hs* CAV1 $_{\beta\text{-ext}}$ , n=22 ROIs were used from two independent experiments. A two-tailed paired t-test was used to calculate p-values, n.s., not significant. (**J**) Representative electron micrograph of isolated and vitrified *h*-caveolae formed by *Hs* CAV1 $_{\beta\text{-ext}}$ expressed in *E. coli*. Scale bar, 50 nm.

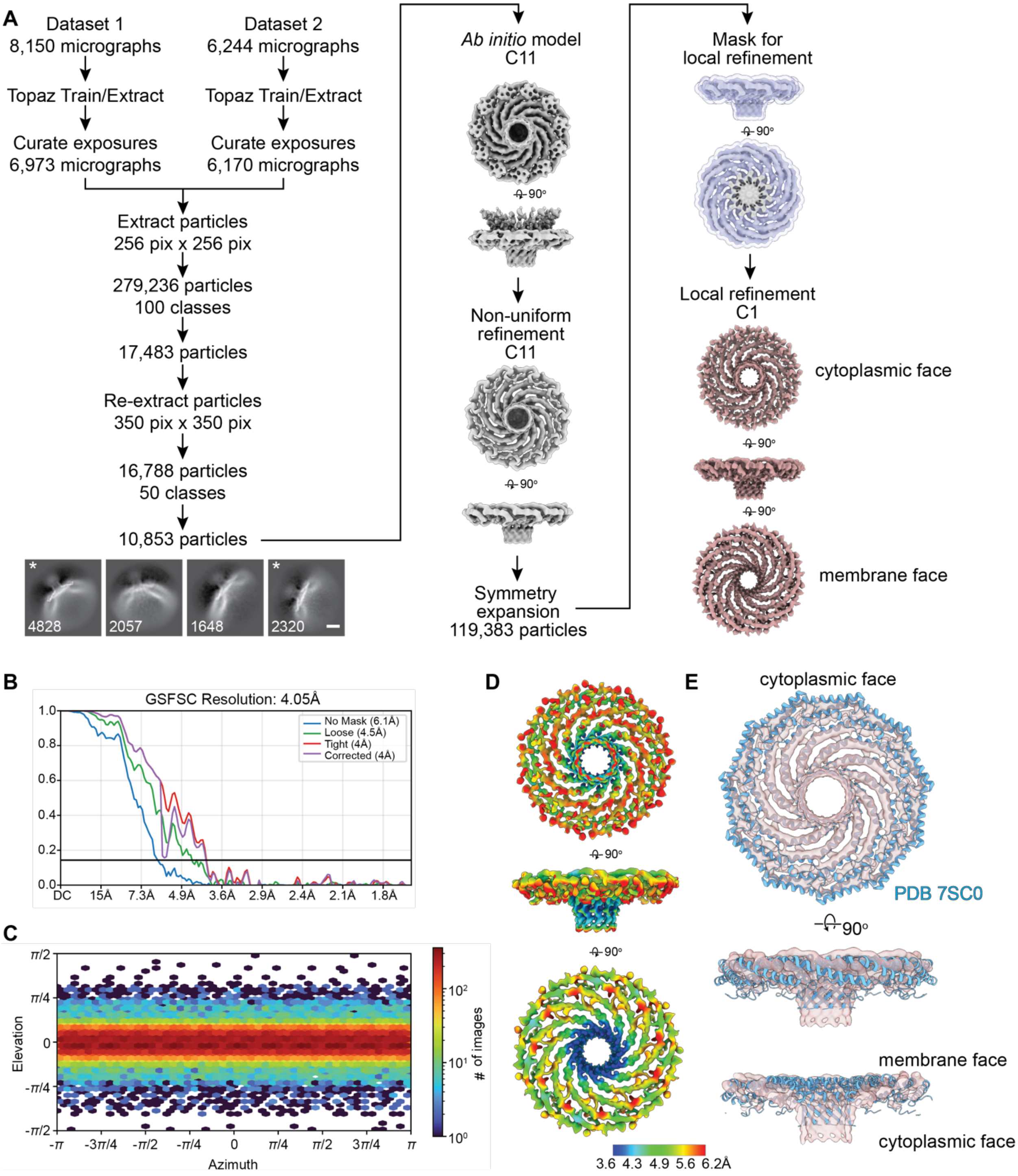

**Figure S11. Single particle cryo-EM data processing of *Hs* CAV1$_{\beta\text{-ext}}$ complexes embedded in membranes.** (**A**) Flowchart depicting the cryo-EM data processing of His$_6$-tagged *Hs* CAV1$_{\beta\text{-ext}}$ embedded in *h*-caveolae membrane carried out using the program cryoSPARC. The number of particles contributing to 2D class averages is noted in the bottom left corner of each average. *, marks 2D averages shown in Figure 5G. All 3D reconstructions are shown in gray in an "*en face"* view and rotated 90° around the x-axis. The mask used for local refinement is shown in transparent light purple. The final 3D reconstruction is shown in rosy brown in an "*en face"* view (cytosolic face), rotated 90° around the x-axis, and rotated a further 90º around the x-axis to the membrane face. Scale bar, 50Å. (**B**) Gold Standard Fourier Shell Correlation (GS-FSC) of final local refinement with no mask (blue line), loose mask (green line), tight mask (red line), and corrected (purple line) done in cryoSPARC. Black horizontal line marks position of Fourier Shell Correlation (FSC) = 0.143. (**C**) Euler angle plot of angles of particle distribution for the final local refinement of the *Hs* CAV1 $_{\beta\text{-ext}}$ complex. (**D**) The *Hs* CAV1$_{\beta\text{-ext}}$ complex map colored by local resolution. The complex is shown in an *en face* view (cytosolic face), rotated 90º around the x-axis to a side view, and rotated a further 90º to the membrane face. (**E**)The model from the EM density map of detergent-solubilized *Hs* CAV1 complex (PDB: 7SC0), shown in blue, placed into the C11 EM non-sharpened density map of the membrane-embedded *Hs* CAV1 $_{\beta\text{-ext}}$ complex, shown in transparent rosy brown. The complex is shown in an *en face* view, rotated 90° around the x-axis to a side view, and a central slice through the side view.

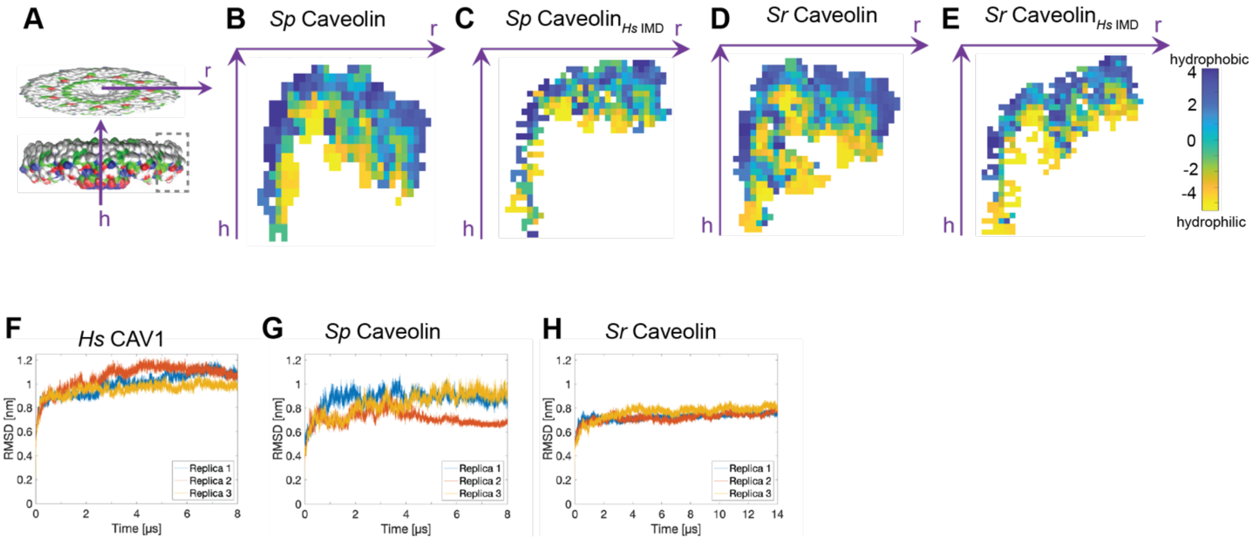

**Figure S12: Hydrophobicity maps of AlphaFold predicted models of *Sp* Caveolin, and *Sr* Caveolin domains swaps and time evolution of caveolin complex during MD simulations.** (**A**) Hydrophobicity mapping protocol. Structure of the *Hs* CAV1 complex (PDB: 7SC0) is colored by residue hydrophobicity: polar green, basic blue, acidic red, and nonpolar white; dashed box indicates the region of the protein-lipid boundary illustrated in panels B-J; h, height; r, radius. (**B-E**) Hydrophobicity maps using the Kyte-Doolittle residue hydrophobicity scale of *Sp* Caveolin (7DN0) (B), AlphaFold model of *Sp* Caveolin$_{Hs\ IMD}$ (C), *Sr* Caveolin (7DN1) (D), and AlphaFold model of *Sp* Caveolin$_{Hs\ IMD}$ (E). h, height; r, radius. (**F-H**) Time evolution of the root means square deviation (RMSD) of the caveolin complexes during the simulation trajectories for *Hs* CAV1 (F), *Sp* Caveolin (G), and *Sr* Caveolin (H). RMSD is shown in units of nanometers as a function of time in units of microseconds for each of the three replica simulations for each variant.

### Supplementary Table 1

| | His$_6$<br>*E. coli* | CAV1-His$_6$<br>*E. coli* | *Sp* Caveolin<br>*E. coli* | *Sr* Caveolin<br>*E. coli* | *Sp* Caveolin$_{W113E}$<br>*E. coli* | *Sr* Caveolin$_{L186E}$<br>*E. coli* | *Sp* Caveolin$_{Hs\ \beta\text{-strand}}$<br>*E. coli* |
|---|---|---|---|---|---|---|---|
| **Data collection and processing** | | | | | | | |
| Microscope | TFS Krios G4i | TFS Krios G4i | TFS Krios G4i | TFS Krios G4i | TFS Krios G4i | TFS Krios G4i | TFS Krios G4i |
| Voltage (kEV) | 300 | 300 | 300 | 300 | 300 | 300 | 300 |
| Camera | Gatan K3 | Gatan K3 | Gatan K3 | Gatan K3 | Gatan K3 | Gatan K3 | Gatan K3 |
| Magnification | 42,000x | 42,000x | 42,000x | 42,000x | 42,000x | 42,000x | 42,000x |
| Pixel size (Å/pixel) | 2.122 | 2.122 | 2.122 | 2.122 | 2.122 | 2.122 | 2.122 |
| Target total electron dose ($e^-/Å^2$) | 120 | 120 | 120 | 120 | 120 | 120 | 120 |
| Defocus range (µm) | -2.0 to -4.0 | -2.0 to -4.0 | -2.0 to -4.0 | -2.0 to -4.0 | -2.0 to -4.0 | -2.0 to -4.0 | -2.0 to -4.0 |
| Energy filter slit width (eV) | 20 | 20 | 20 | 20 | 20 | 20 | 20 |
| Data used | 10 | 10 | 10 | 10 | 10 | 10 | 6 |

| | *Sr* Caveolin$_{Hs\ \beta\text{-strand}}$<br>*E. coli* | *Sp* Caveolin$_{Hs\ IMD}$<br>*E. coli* | *Sr* Caveolin$_{Hs\ IMD}$<br>*E. coli* | *Hs* CAV1$_{E140A}$<br>*E. coli* | *Hs* CAV1$_{Sp\ \beta\text{-strand}}$<br>*E. coli* | *Hs* CAV1$_{Sr\ \beta\text{-strand}}$<br>*E. coli* | *Hs* CAV1$_{Sp\ IMD}$<br>*E. coli* |
|---|---|---|---|---|---|---|---|
| **Data collection and processing** | | | | | | | |
| Microscope | TFS Krios G4i | TFS Krios G4i | TFS Krios G4i | TFS Krios G4i | TFS Krios G4i | TFS Krios G4i | TFS Krios G4i |
| Voltage (kEV) | 300 | 300 | 300 | 300 | 300 | 300 | 300 |
| Camera | Gatan K3 | Gatan K3 | Gatan K3 | Gatan K3 | Gatan K3 | Gatan K3 | Gatan K3 |
| Magnification | 42,000x | 42,000x | 42,000x | 42,000x | 42,000x | 42,000x | 42,000x |
| Pixel size (Å/pixel) | 2.122 | 2.122 | 2.122 | 2.122 | 2.122 | 2.122 | 2.122 |
| Target total electron dose ($e^-/Å^2$) | 120 | 120 | 120 | 120 | 120 | 120 | 120 |
| Defocus range (µm) | -2.0 to -4.0 | -2.0 to -4.0 | -2.0 to -4.0 | -2.0 to -4.0 | -2.0 to -4.0 | -2.0 to -4.0 | -2.0 to -4.0 |
| Energy filter slit width (eV) | 20 | 20 | 20 | 20 | 20 | 20 | 20 |
| Data used | 8 | 10 | 10 | 10 | 6 | 9 | 10 |

| | *Hs* CAV1$_{Sr\ IMD}$<br>*E. coli* | mVenus-His$_{10}$<br>*E. coli* | *Hs* CAV1-mVenus-His$_{10}$<br>*E. coli* | Purified *Hs* CAV1-mVenus-His$_{10}$<br>*h*-caveolae | *Hs* CAV1$_{\beta\text{-ext}}$<br>*E. coli* |
|---|---|---|---|---|---|
| **Data collection and processing** | | | | | |
| Microscope | TFS Krios G4i | TFS Krios G4i | TFS Krios G4i | TFS Krios G4i | TFS Krios G4i |
| Voltage (kEV) | 300 | 300 | 300 | 300 | 300 |
| Camera | Gatan K3 | Gatan K3 | Gatan K3 | Gatan K3 | Gatan K3 |
| Magnification | 42,000x | 42,000x | 42,000x | 42,000x | 42,000x |
| Pixel size (Å/pixel) | 2.122 | 2.122 | 2.122 | 2.122 | 2.122 |
| Target total electron dose ($e^-/Å^2$) | 120 | 120 | 120 | 100 | 120 |
| Defocus range (µm) | -2.0 to -4.0 | -2.0 to -4.0 | -2.0 to -4.0 | -4.0 to -5.0 | -2.0 to -4.0 |
| Energy filter slit width (eV) | 20 | 20 | 20 | 20 | 20 |
| Data used | 10 | 14 | 10 | 30 | 10 |

**Supplementary Table 2. Primers used to generate mammalian expression constructs.**

| **Plasmid name** | **Forward primer, 5'-3'** | **Reverse primer, 5'-3'** |
|---|---|---|
| *Sr* Caveolin | GCGGCCGGTACCATGAGCTACCAC | GCGGCCCCGCGGTTGTCCTTCAGTTCCTTGTGG |
| *Sr* Caveolin $_{Hs\ \mathrm{IMD}}$ | same as above | same as above |
| *Sr* Caveolin $_{\mathrm{L186E}}$ | CCAGCTGATGTTCCTGGAGATCGCCAGCCTGGCC | GGCCAGGCTGGCGATCTCCAGGAACATCAGCTGG |
| *Sr* Caveolin $_{Hs\ \beta\text{-strand}}$ | CTGGCTCTGAGCAACGTCCGCATCAACTTGCAGAAAGAAATAAAGCTTGGGCCCGAAC | GTTCGGGCCCAAGCTTTATTTCTTTCTGCAAGTTGATGCGGACGTTGCTCAGAGCCAG |
| *Sp* Caveolin | GCGGCCGGTACCATGGAACTGATCCATCCTG | GCGGCCCCGCGGTTCACCTGGCTGGTCTTAACATCAGAGC |
| *Sp* Caveolin $_{Hs\ \mathrm{IMD}}$ | same as above | same as above |
| *Sp* Caveolin $_{\mathrm{W113E}}$ | CCTATCATGAAACTGTACTACGTGGAGCTGCGGGTGTTCAACC | GGTTGAACACCCGCAGCTCCACGTAGTACAGTTTCATGATAGG |
| *Sp* Caveolin $_{Hs\ \beta\text{-strand}}$ | CTGATCCTGAGCGGCGTCCGCATCAACTTGCAGAAAGAAATAAAGCTTGGGCCCGAAC | GTTCGGGCCCAAGCTTTATTTCTTTCTGCAAGTTGATGCGGACGCCGCTCAGGATCAG |
| *Hs* CAV1 $_{\mathrm{E140A}}$ | GAGCTTCCTGATTGCCATTCAGTGCATCAGCCG | CGGCTGATGCACTGAATGGCAATCAGGAAGCTC |
| *Hs* CAV1 $_{Sp\ \beta\text{-strand}}$ | CTGTTGGGAAAATATTCAGCAATATCAAGGGCCAGTTCAAGATGAACAGCTCTGATGTTAAGACCAGCCAGGTGAAGCTTGGGCCCGAAC | GTTCGGGCCCAAGCTTCACCTGGCTGGTCTTAACATCAGAGCTGTTCATCTTGAACTGGCCCTTGATATTGCTGAATATTTTCCCAACAG |
| *Hs* CAV1 $_{Sr\ \beta\text{-strand}}$ | GTTGGGAAAATATTCAGCAATATCCAGGTCAGACTGGGCATGGAAGGCAAGGCCCGCCACAAGG | CCTTGTGGCGGGCCTTGCCTTCCATGCCCAGTCTGACCTGGATATTGCTGAATATTTTCCCAAC |
| *Hs* CAV1$_{\beta\text{-ext}}$ | CCGCATCAACTTGCAGAAAGAAATAAAGGCCCGCCACAAGGAACTGAAGGACAAGCTTGGGCCCG | CGGGCCCAAGCTTGTCCTTCAGTTCCTTGTGGCGGGCCTTTATTTCTTTCTGCAAGTTGATGCGG |

**Supplemental Table 3**

| | *H. sapiens* $CAV1_{\beta\text{-ext}}$ in *h*-caveolae<br>EMDB-XXXXX<br>PDB-XXXX |
|---|---|
| **Data collection and processing** | |
| Magnification | 105,000x |
| Voltage (kEV) | 300 |
| Total electron exposure ($e^{-}/Å^2$) | 50 |
| Defocus range (μm) | -0.5 to -3.0 |
| Pixel size (Å/pixel) | 0.83 |
| Symmetry imposed | C11 |
| Initial particles (number) | 279,236 |
| Final particles (number) | 10,853 |
| Map resolution (Å) | 4.05 |
| FSC threshold | 0.143 |
| Map resolution range (Å) | 3.6 to 8.5 |
| **Refinement** | |
| Initial model used (PDB code) | 7SC0 |
| Model resolution (Å) | 4.2 |
| Map sharpening *B* factor ($Å^2$) | 163.8 |
| Model composition | |
| Non-hydrogen atoms | 9,042 |
| Protein residues | 1,100 |
| Ligands | None |
| *B* factors ($Å^2$) | |
| Protein | -130 |
| Ligand | None |
| R.m.s. deviations | |
| Bond lengths (Å) | 0.004 |
| Bond angles (º) | 0.820 |
| Validation | |
| MolProbity score | 2.33 |
| Clash score | 26.94 |
| Poor rotamers (%) | 0.00 |
| Ramachandran plot | |
| Favored (%) | 93.88 |
| Allowed (%) | 6.12 |
| Disallowed (%) | 0.00 |
